# Thermal proteome profiling identifies glucose binding proteins involved in metabolic disease

**DOI:** 10.1101/2025.05.06.651509

**Authors:** Lindsey M. Meservey, Ian D. Ferguson, Vivian B. Tien, Luca Ducoli, Weili Miao, David M. Lu, Suhas Srinivasan, Vanessa Lopez-Pajares, Douglas F. Porter, Zhongwen Cao, Yisheng Wang, Paul A. Khavari

## Abstract

Recent work has shown that glucose can directly bind non-enzymatic proteins to modulate their function. We sought to build on this work by adapting thermal proteome profiling screens to uncover novel glucose interactors involved in metabolism. Proteome integral solubility alternation (PISA) profiling nominated 23 proteins with glucose-induced thermal solubility, including two proteins strongly implicated in metabolic disease, TSC22D4 and IGF2BP2. First, we characterized glucose binding of TSC22D4, an intrinsically disordered leucine zipper protein strongly implicated in hepatic steatosis. MST confirmed direct glucose-protein interaction of TSC22D4. UV-crosslinking mass spectrometry identified putative glucose binding at the C-terminal leucine zipper. Mutating isoleucine 322 to tryptophan (I322W) abolishes glucose binding. Crosslinking-MS and chemo-proteomic experiments suggest that glucose increases accessibility of the leucine zipper region, resulting in intra-protein contacts between C-terminal zipper domain and N-terminal intrinsically-disordered domain. TSC22D4 associates with fatty acid metabolism machinery proteins in response to high glucose conditions, suggesting a possible role for TSC22D4 in altering fatty acid metabolism. Next, we characterized glucose-binding in IGF2BP2, an RNA-binding protein essential for beta cell insulin secretion. We first confirmed direct IGF2BP2-glucose interaction and created glucose-binding mutant Y40A based on computational docking predictions. IGF2BP2^Y40A^ exhibited a dominant negative impact on proliferation and insulin secretion in MIN6-6 beta cells. Glucose increased IGF2BP2 binding to client mRNAs Igf2 and Pdx1, which has previously been demonstrated to mediate its impact on insulin secretion. Thus, our data suggests that IGF2BP2 directly binds glucose to enhance insulin secretion.

## Introduction

Protein-metabolite interactions modulate protein function.^1^ Pioneering work showed transcription factor LXR directly binds glucose and glucose-6-phosphate to facilitate binding to coactivator recruitment and target gene transcription in liver cells.^2^ New findings suggest glucose engages directly with a broader range of nucleotide-binding proteins to modulate their multimerization, nucleotide binding, and cellular localization.^3–6^ Glucose gradients formed during stratified epithelial differentiation lead to induction of binding events in key drivers of differentiation.^3,4^ Specifically, glucose binding leads to dimerization and genome targeting of IRF6, a pioneer transcription factor in keratinocyte differentiation, which drives induction of differentiation related genes.^3^ Glucose accumulation and binding leads to de-dimerization and re-localization of DDX21, which alters RNA-splicing to drive differentiation.^4^

Importantly, these glucose binding proteins were identified by affinity purification mass spectrometry strategies using dextran column or biotin-glucose probes, which both require modified glucose derivatives. Another rapidly developing approach to study small protein-ligand interactions is thermal proteome profiling (TPP), which measures binding-induced changes in thermal solubility with unmodified metabolites.^1,7,8^ In this work, we apply TPP methods to identify new glucose binding proteins and characterize the effect of glucose binding on protein function for two new glucose targets, TSC22D4 and IGF2BP2, which have established genetic linkage to fatty liver disease and diabetes, respectively.^9–13^

## Results

### Thermal profiling proteomics nominates glucose-binding proteins

We first sought to utilize TPP techniques to expand upon existing research identifying glucose binding proteins. Traditionally, TPP has been used to discover high affinity ligand-protein interactions such as small molecule inhibitors.^14^ Recently, TPP of small metabolites was performed in *E. coli* lysate using supraphysiological osmolyte concentrations.^15^ We hypothesized that TPP at physiological osmolyte concentrations could uncover functional hexose-protein interactions without leading to global proteome condensation as has previously been observed.^16^

We opted to perform proteome integral solubility alternation (PISA) assays, in which soluble fractions from each temperature are pooled, to allow for multiple treatment replicates run in a single TMT experiment (Fig 1A).^7,17^ We validated that glucose-induced solubility of HK1 was visible in a PISA-western and saw slight solubilization of IRF6, a known glucose binder with a K_d_ of 144 μM, illustrating that some glucose binders might be less amenable to thermal shift assays (Fig S1A-D).^3^ Total proteome solubilization was unchanged by glucose, and fold change of glucose induced stabilization of HK1 was similar to that of ATP-induced Actin stabilization (Fig S1E-G).

**Figure 1.**
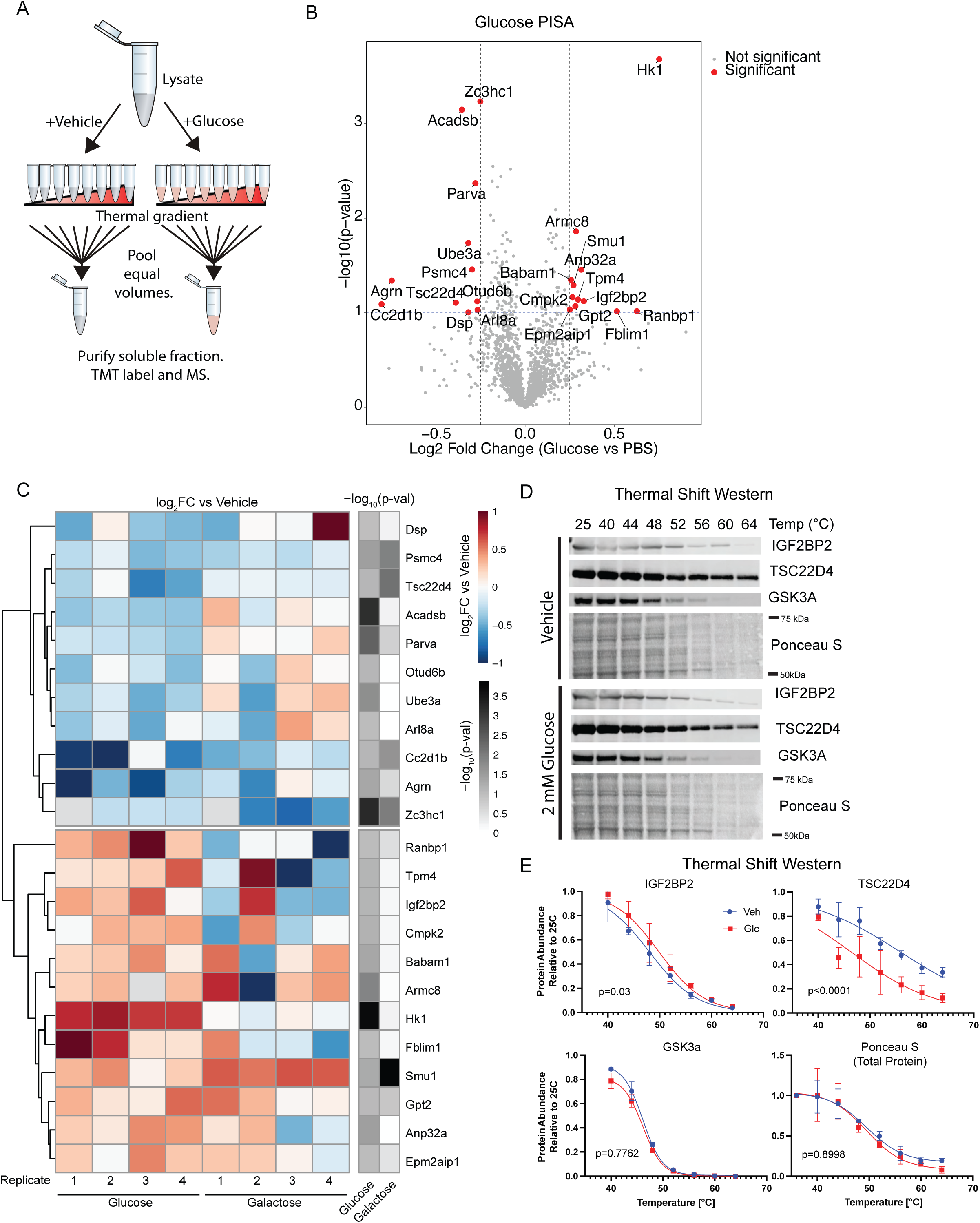
Thermal profiling proteomics nominates glucose-binding proteins at physiologic glucose concentrations. **A**) Schematic of PISA experiment **B**) Proteome integral solution assays (PISA) performed in lysates extracted from 3T3-L1 pre-adipocytes **C**) Log_2_ fold thermal stability changes and p-value observed in PISA with glucose and galactose treatments for nominated glucose binders **D**) Thermal solubility assay western profile for human homologs of nominated glucose-binding proteins IGF2BP2 and TSC22D4 with GSK3A and total protein stain (Ponceau) as control in HEK293T lysates, quantified in **E**

We performed PISA assays in 3T3-L1 mouse pre-adipocyte lysates, reasoning that glucose-binding proteins could represent critical regulators of adipocyte differentiation, a process that is tightly coupled to glucose availability^18^ and characterized by increased intracellular glucose concentrations.^3^ We treated 3T3-L1 lysate with vehicle, glucose, or stereoisomer galactose. Generally, galactose thermal solubility shifts were only weakly correlated with glucose after normalization to vehicle treatment, suggesting that glucose exhibits structure-specific protein solubilization (Fig S1H). We identified 23 proteins whose solubility significantly shifted with glucose treatment (Fig 1B-C). Glucose-binding proteins enriched for nucleotide-binding proteins, consistent with previous screens for glucose-binding proteins (Fig S1I)^3–5^. Among the novel glucose-binding proteins identified by our studies, TSC22D4 and IGF2BP2 stood out in their connection to metabolic disease.^9–12^ TSC22D4 has been connected with hepatic insulin signaling, lipogenesis, and oxygen consumption^9,13,19^. IGF2BP2 has been genetically linked with global insulin sensitivity and pancreatic insulin secretion^20–22^. TSC22D4 was de-solubilized by both glucose and galactose, and IGF2BP2 was solubilized by glucose but not galactose (Fig 1C). We validated that the human orthologs of TSC22D4 and IGF2BP2 had shifted thermal solubility in human cells by thermal shift western (Fig 1D-E).

### TSC22D4 binds glucose at TSC box

First, we validated that TSC22D4 directly binds glucose using microscale thermophoresis (MST), calculating K_d_ ≈ 464 μM (Fig S2A). Importantly, TSC22D4 does not strongly bind glucose-6-phosphate at physiological levels (Fig S2A). To identify residues involved in glucose binding, we employed UV-crosslinking mass spectrometry and obtained a high-confidence peptide, IEQAMDLVK, crosslinked to glucose (Fig. 2A-B, Fig. S2B-D). Mutation I322W ablated glucose binding, while I322P, I322G and triple mutant I322G/E323G/Q324G did not alter glucose binding (Fig. 2C, Fig S2E-F,H). Importantly, TSC22D4^I322W^ was still able to dimerize, suggesting proper folding structure (Fig. S2G-H). We next applied a recently developed approach for assessing lysine accessibility using formaldehyde-based chemoproteomics^23^, and found increased labeling of a C-terminal lysine at position 345 after incubation with glucose (Fig 2D), suggesting increased accessibility of the C-terminal region of the protein. To further understand how glucose modulates TSC22D4 monomeric conformation, we employed BS3 crosslinking on TSC22D4 incubated with glucose, isolating the monomeric band for mass spectrometry (Fig S2I). Glucose incubation induced new long-range contacts between the C-terminus of the protein in the TSC Box to the N-terminus of the protein (Fig 2E), suggesting that glucose alters TSC box contacts, in agreement with the hypothesis that glucose increases accessibility the C-terminus of TSC22D4. Glucose had no significant effect on TSC22D4 dimerization in MST experiments (Fig 2F), although it did shift TSC22D4 to smaller complexes in fast-protein liquid chromatography (Fig 2G, Suppl S2J).

**Figure 2.**
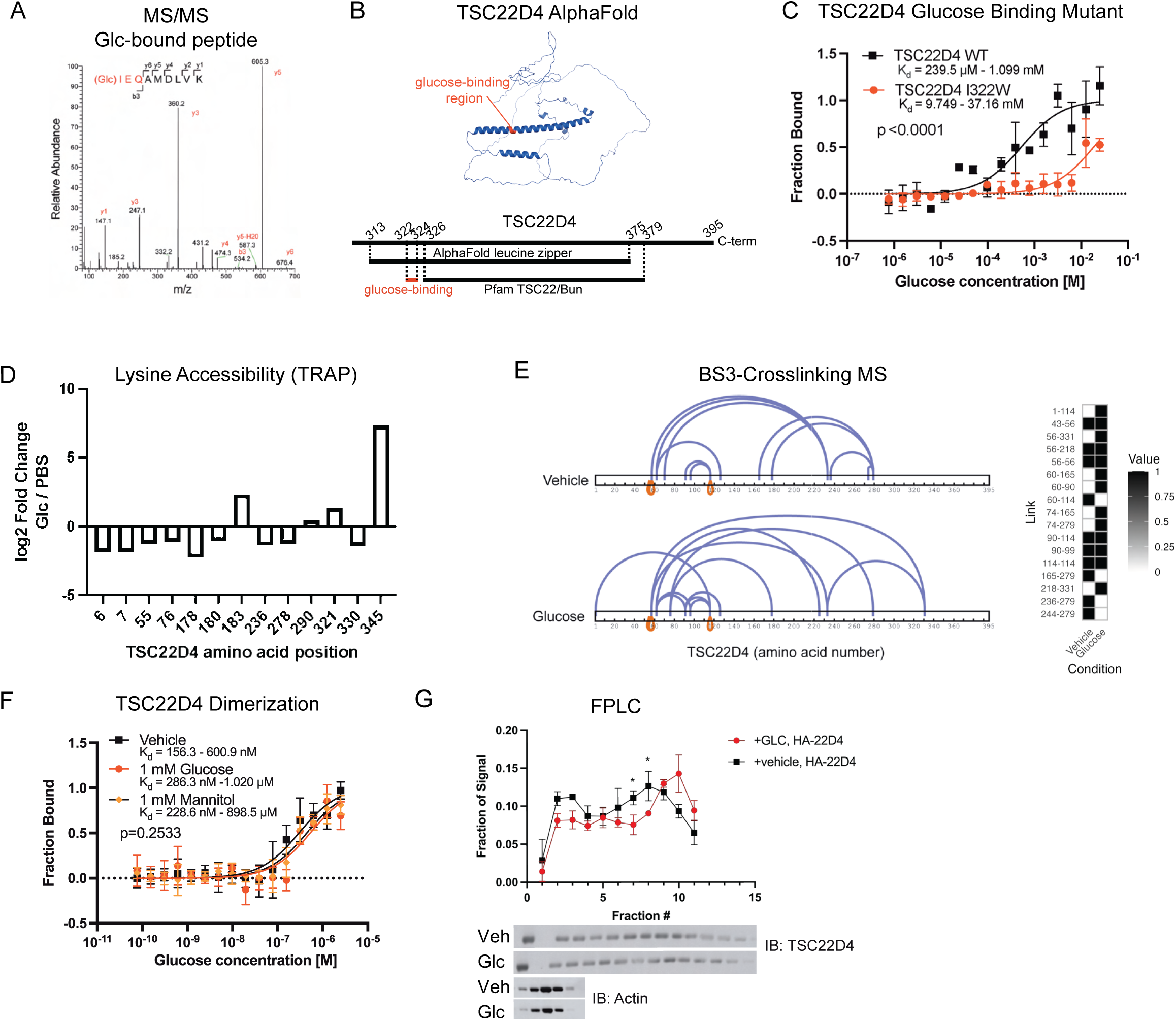
TSC22D4 binds glucose at TSC box to increase C-terminal accessibility. **A**) MS/MS for UVC bound peptide IEQAMDLVK. **B**) AlphaFold structure^36–38^ of TSC22D4 annotated with glucose-binding region in red, and comparative localization of structurally predicted leucine zipper with annotated Pfam TSC22/Bun domain **C**) MST analysis for TSC22D4-I322W showing decreased binding to glucose. **D**) Lysine accessibility chemoproteomics showing increased labeling of lysine 345 at the C-terminus of TSC22D4. **E**) Intra-protein BS3 crosslinking showing induced contacts between C-terminus and N-terminus under glucose treatment. **F**) MST dimerization analysis using labeled His-TSC22D4 and un-labeled TSC22D4 showing that neither glucose nor mannitol significantly impacts TSC22D4 dimerization. **G**) FPLC-western for HEK293T lysates treated with glucose or vehicle shows TSC22D4 distribution shift to smaller protein complexes

### Glucose modulates TSC22D4 interactome

As TSC22D4 is an annotated chromatin binder and transcriptional repressor, and other transcription factors have shown to alter DNA association upon glucose binding, we first tested whether glucose alters TSC22D4 chromatin association. In vitro, glucose decreased TSC22D4 affinity toward DNA in a non sequence-specific manner (Fig S3A-B). We performed Cut&Run on 3T3-L1s expressing HA-TSC22D4, but despite technical success, we were unable to see any robust TSC22D4 genomic binding regions by Cut&Run in either glucose condition (Fig S3C). We performed ChIP but were unable to confirm any chromatin binding of TSC22D4, despite experimental success with the same antibody (Fig S3D-E). Our lack of evidence showing TSC22D4 interacting with chromatin led us to entertain alternative hypotheses.

Based on the glucose-dependent shift in protein complexing together with literature reports that TSC22D4 scaffolds protein-protein interactions, we hypothesized that glucose modulates TSC22D4 interactome.^9,24^ We performed affinity purification (AP)-MS in 3T3-L1 cells to map the glucose-dependent TSC22D4-interactome. We found that crosslinking in-lysate with BS3 improved pulldown of tagged and endogenous TSC22D4, suggesting crosslinking improved recovery of dimeric TSC22D4 (Fig. 3A). Therefore, we proceeded with BS3-crosslinking for AP-MS, treating 3T3-L1 cells for 1.5 hours with either glucose, 3-MG, or vehicle. Since 3-MG does not enter metabolic pathways, its close alignment with glucose in altering the TSC22D4 interactome indicates that the observed shifts are primarily due to non-metabolic effects, such as osmolarity or protein-metabolite binding (Fig. S3A). Shared interactors across all conditions included NRBP2, a previously profiled TSC22D4 interactor (Fig. 2B, S4B-D).^25^ TSC22D4 interactions with actin and cytoskeleton proteins were enriched in the vehicle treated cells, while interactions with proteasome and vesicle transport proteins were increased in glucose conditions (Fig. 3C). Clustering proteins enriched >2-fold in glucose conditions using Markov Clustering^26^ on StringDB^27^, we again saw an enrichment for vesicle transport machinery (Fig 3D). We were surprised to observed a glucose-dependent interaction with respiratory chain and fatty acid metabolism machinery, including key regulators of fatty acid synthesis ACACA and AACS (Fig 3D-E). We performed mitochondrial isolation to find TSC22D4 associates with mitochondria, suggesting a direct interaction between TSC22D4 and mitochondrial metabolic enzymes (Fig 3F).

**Figure 3.**
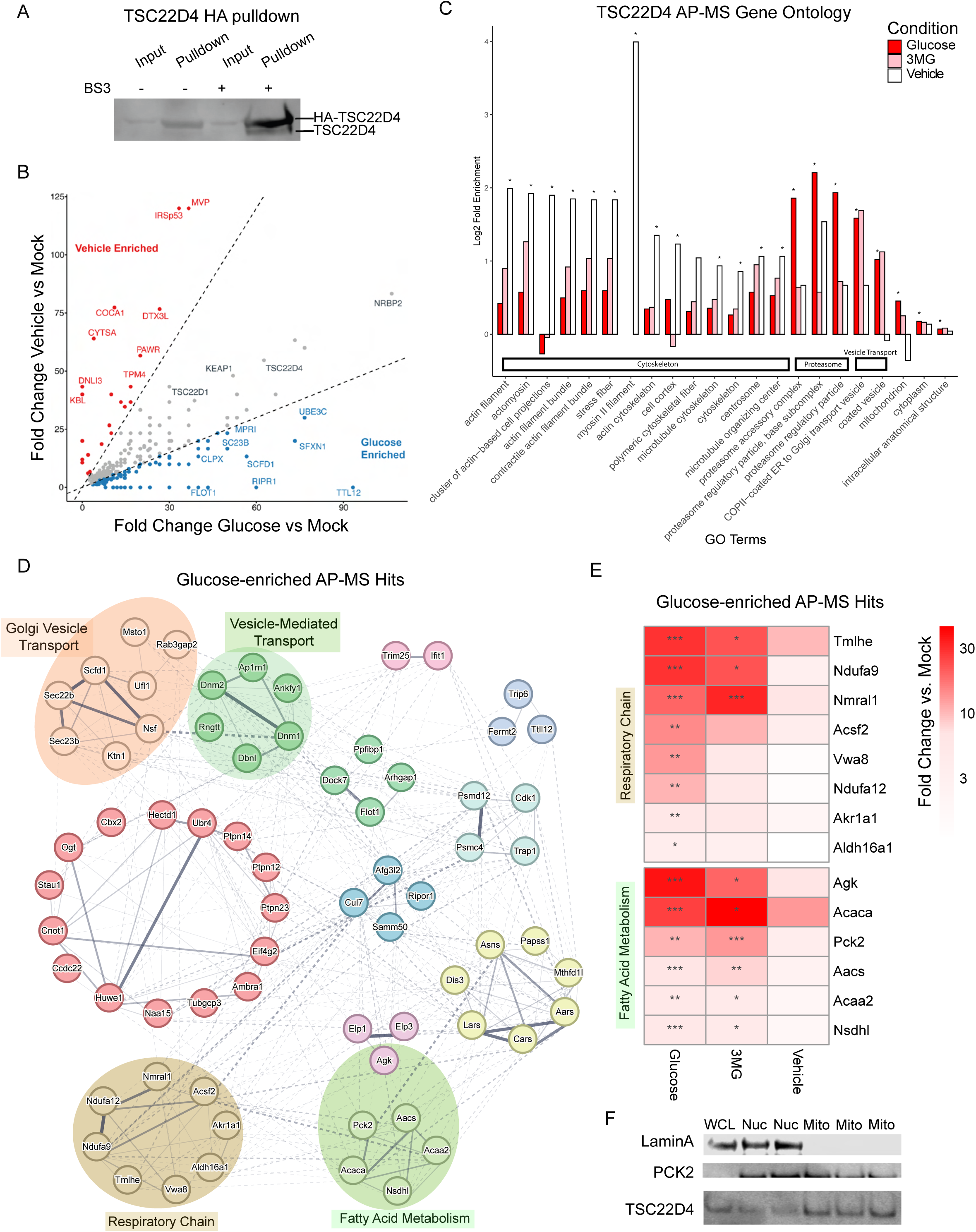
Glucose modulates TSC22D4 interactome. **A**) Pulldown of HA-TSC22D4 shows increased efficiency of dimer pulldown under BS3 crosslinking vs native conditions **B**) Scatterplot comparing fold change of prey in glucose and vehicle conditions. Dotted lines indicate 2-fold enrichment. **C**) Gene ontology analysis of proteins enriched (SAINT >= 0.8) in glucose, 3MG, and vehicle conditions. **D**) Proteins enriched >2-fold in both glucose and 3MG condition vs vehicle (with SAINT score > 0.7 for either glucose or 3MG) were clustered by Markov Clustering (MCL) algorithm^26^ on StringDB^27^ (inflation parameter = 2.3). **E**) Heatmap of glucose- and 3MG-enriched AP-MS hits in respiratory chain and fatty acid metabolism clusters. * SAINT > 0.65, ** SAINT > 0.80, *** SAINT > 0.95. **F**) Western blot shows TSC22D4 associates with mitochondria in 3T3-L1. WCL = whole cell lysate, Nuc = Crude Nuclear Fraction, Mito = Purified Mitochondrial Fraction.

### IGF2BP2 binds glucose at RRM1 domain to modulate insulin secretion

We next sought to validate glucose binding to IGF2BP2, another putative target from the PISA screen (Fig 1B-E). IGF2BP2 has been connected to type 2 diabetes by GWAS studies from multiple populations.^28–30^ IGF2BP2 loss improves insulin sensitivity and protects against diet-induced obesity by altering stability and translational efficiency of client m^6^A-modified mRNAs.^12,31^ We purified IGF2BP2, performed MST, and found a K_d_ for glucose of 8.2 μM (Fig 4A). Glucose did not impact IGF2BP2 RNA binding or dimerization in vitro (Fig S6A-B). Boltz-2 nominated IGF2BP2’s RRM1 domain as a glucose-binding site at well-conserved residue (Fig 4B, Fig S6C-D).^32^ This localization was supported by experimental data in IGF2BP3, a homolog of IGF2BP2 previously nominated as a glucose-binding protein (Fig S7A-H). IGF2BP3^Y39A^ showed abolished glucose binding without large decrease in RNA binding in cell culture (Fig S7I-L). We confirmed experimentally that IGF2BP2^Y40A^ does not bind glucose (Fig 4C).

**Figure 4.**
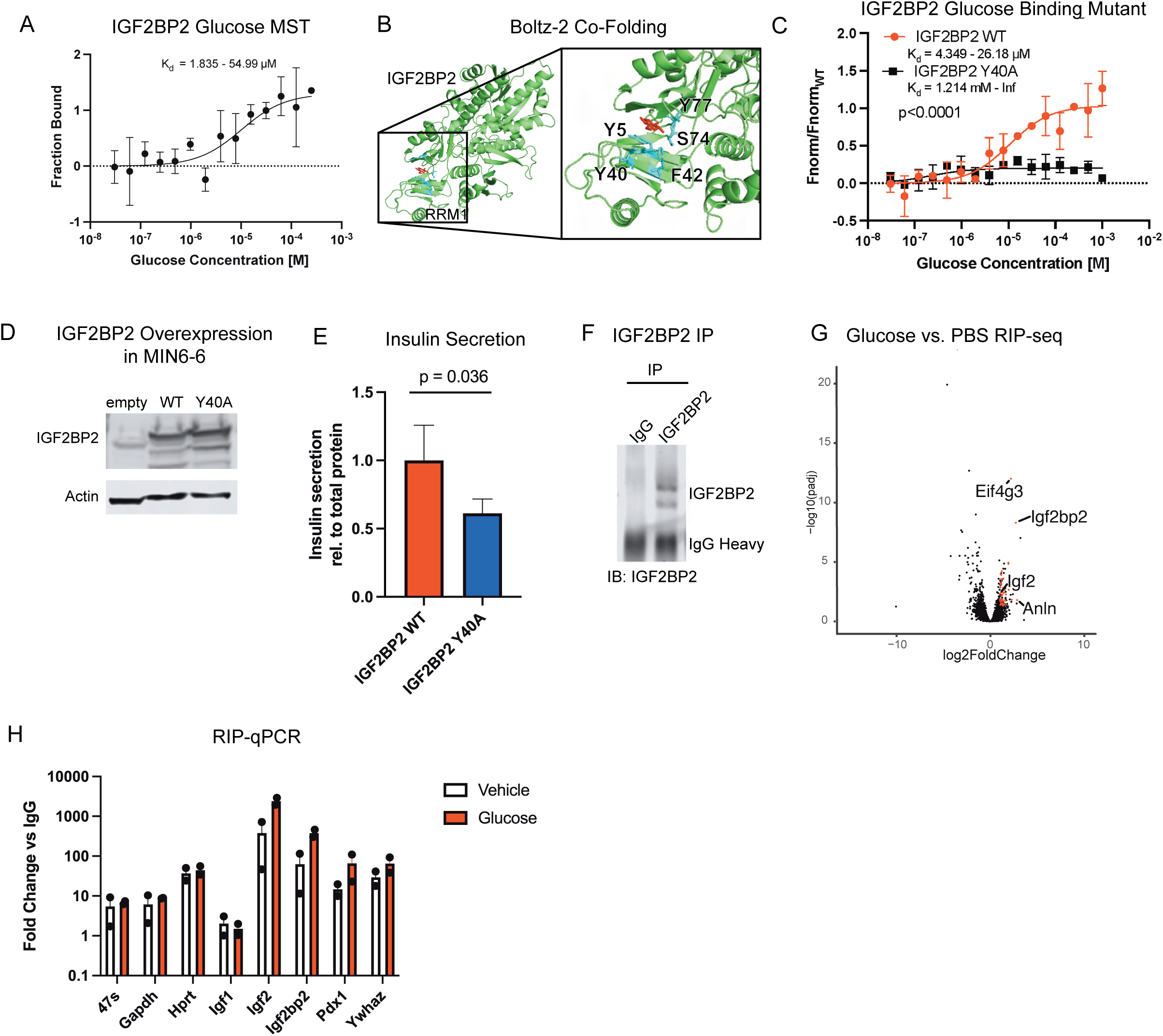
IGF2BP2 binds glucose at RRM1 domain to modulate insulin secretion. **A**) MST analysis of IGF2BP2 shows 12 μM affinity for glucose. **B**) Boltz-2 co-folding of IGF2BP2 and glucose predicting interaction within RRM1 domain **C**) MST analysis shows Y40A mutant ablates IGF2BP2-glucose binding **D**) Western blot analysis showing overexpression of IGF2BP2 WT or Y40A protein in MIN6-6 cells. **E**) Luciferase-based insulin secretion assay in MIN6-6 cells showing dominant negative effect of Y40A variant overexpression. **F**) IP of native IGF2BP2 from MIN6-6 cells for RIP-qPCR. **G**) Volcano plot of RIP-seq, with red dots showing enrichment in both IGF2BP2 vs. IgG and glucose vs. PBS. **H**) RIP-qPCR in MIN6-6 cells starved of glucose for 2h and treated with glucose or vehicle for 1.5 hours.

Next, we sought to study the effect of IGF2BP2 glucose binding on IGF2BP2 function in pancreatic beta cell insulin secretion. IGF2BP2 has been shown to increase insulin secretion by stabilizing *Pdx1* mRNA, which encodes a master transcriptional regulator of the insulin production pathway.^12^ We tested the impact of IGF2BP2-glucose interaction on insulin secretion in MIN6-6, an immortalized mouse pancreatic beta cell line with luciferase tagged insulin.^33,34^ We overexpressed IGF2BP2^WT^ or IGF2BP2^Y40A^ in MIN6-6 and found that IGF2BP2^Y40A^ had a dominant negative effect on cell proliferation and insulin secretion (Fig 4D-E, Fig S6E). Because IGF2BP2 regulates insulin secretion by stabilizing client RNAs, we performed RNA immunoprecipitation (RIP)-seq to characterize the IGF2BP2 glucose-dependent targetome (Fig 4F,G). We found many differentially enriched target mRNAs, including *Eif4g3*, *Igf2bp2*, and *Igf2*. We further validated these targets by RIP-qPCR (Fig 4H, note log10 scale).

## Discussion

Here we apply TPP approaches to identify new glucose binding proteins and characterize two novel glucose-binding proteins, TSC22D4 and IGF2BP2, genetically linked to metabolic disease. TSC22D4 binds glucose near its C-terminal TSC box, leading to increased accessibility of the C-terminal region. Mutant TSC22D4^I322W^ ablates glucose binding. Glucose binding alters TSC22D4-DNA binding affinity in vitro; however, we failed to detect any chromatin binding events in cells. Glucose binding shifts the TSC22D4 interactome away from cytoskeletal proteins and toward vesicle transport, respiratory chain, and fatty acid metabolism machinery. Future experiments should test whether TSC22D4 localizes to the mitochondria in a glucose-dependent manner and whether TSC22D4 alters fatty acid metabolism.

IGF2BP2 binds glucose at its RRM1 domain, and mutant IGF2BP2^Y40A^ showed ablated glucose binding. In pancreatic beta cells, IGF2BP2^Y40A^ overexpression had a dominant negative effect on proliferation and insulin secretion. Glucose increased IGF2BP2 association with client mRNAs *IGF2*, *PDX1*, and *IGF2BP2*, which are essential for beta cell maintenance and insulin secretion. Together, these results support the hypothesis that glucose binding increases IGF2BP2 binding to client mRNAs, whose translation is essential for proper insulin secretion. Future efforts should confirm whether these binding events are directly dependent on IGF2BP2-glucose interaction and how IGF2BP2-glucose interaction impacts RNA metabolism and translational efficiency.

Previous work on GBPs has shown glucose rewires GBP function through altered oligomerization^3–5,35^. However, neither TSC22D4 nor IGF2BP2 show altered oligomerization upon treatment with glucose. However, TSC22D4 exhibited vastly altered protein complexing upon glucose treatment. This suggests that glucose may rewire protein interactions independent of homo-oligomerization.

We also show that IGF2BP2 homolog IGF2BP3 is a glucose-binding protein and characterize glucose-binding mutant IGF2BP3^Y39A^. Previous work has shown homologs DDX21 and DDX50 bind glucose at homologous site.^4,35^ Thus, there appears to be a clear structural component to glucose binding. However, as global characterized glucose-binding sites are incredibly diverse and share no obvious structural homology, and primary sequence alignment fails to give a clear primary sequence motif, which structural features dictate glucose (or other glycolytic intermediate) binding remains an outstanding question. As we have experienced Boltz-2 replicate experimental data for glucose-binding sites, we are optimistic that AI technologies originally trained for protein-drug interaction may be repurposed to nominate the structural features essential for small molecule binding.

This work suggests that, in addition to canonical nutrient signaling pathways, cells employ a complementary layer of regulation in which proteins directly bind glucose to modulate their functions. Here, we elucidate the role of two such proteins in detail, providing mechanistic insights into how direct glucose binding can alter cellular processes. Beyond these examples, our study also nominates tens of previously uncharacterized candidate glucose-binding proteins, adding to the growing list of proteins proposed to directly sense this metabolite. Taken together, these findings support the emerging view that nutrient sensing is not restricted to well-established signaling cascades but is also mediated by a broad and intricate network of proteins that can directly recognize glucose and other small metabolites. Such a distributed sensing system may enable cells to integrate diverse metabolic cues with high specificity and adaptability.

In sum, we extend previous work on GBPs in epidermal differentiation into two phenotypic systems where glucose dysregulation is a common mechanism and pathology of disease—insulin secretion and adipogenesis. This work suggests that, in addition to nutrient sensors with well-established roles in metabolic disease (e.g., AMPK, mTOR, ChREBP), direct glucose sensors contribute to metabolic disease. Further work is required to fully understand the mechanisms by which glucose binding alters TSC22D4 and IGF2BP2 function in adipogenesis and insulin secretion.

## Acknowledgements

We thank all members of the Khavari laboratory for their valuable input, feedback, and support throughout the course of this work. We are also grateful to Dr. Seung Kim and Dr. Jonathan Long for their insightful discussions and generous sharing of expertise, which significantly enriched this study.

## Methods

### Cell culture

Lenti-X™ 293T (Takara Bio #632180) and 3T3-L1 cells were grown in DMEM + 10% FBS + PenStrep. HIT-T15 were grown in RPMI + 10% FBS + PenStrep. MIN6-6 cells were grown in DMEM + 10% FBS + PenStrep + 10 mM HEPES (Thermo #15-630-080) + 55 uM 2-Mercaptoethanol (Thermo #21985023). MIN6 cells were grown in DMEM + 15% FBS + PenStrep + 10 mM HEPES (Thermo #15-630-080) + 55 uM 2-Mercaptoethanol (Thermo #21985023). Beta-TC-6 were grown in DMEM + 15% FBS + 1% PenStrep. Cells were regularly tested for Mycoplasma contamination.

### Proteome Integral Solubility Alteration

3T3-L1 cells were grown to confluency in DMEM media and then starved overnight in 0g/L Glc DMEM+D-FBS. Cells collected by trypsinization (with quenching by 0g/L DMEM). Cells were washed 3x with ice-cold D-PBS. Pellet lysed in PISA-Lysis buffer (50 mM Tris-HCl pH 7.5, 150 mM NaCl, 1.5 mM MgCl2, 0.5% NP-40) + PIC (Sigma), incubated on ice for 15 min, and cleared with 20,000g spin at 4°C for 15 min. Supernatant was collected and protein quantified by BCA assay (Pierce). Lysate was diluted to 2.2mg/mL and split across 16 tubes and treated with 350 μM hexose. Samples briefly spun down in microcentrifuge and incubated at room temperature for 30 minutes without rotation or shaking, followed by separating each sample into 8 tubes for heating in thermal cycler at 44, 46, 48, 50, 52, 54, 56, 58 °C. After heating allowed to return to RT for 5 minutes. Equal volumes of each temperature were pooled prior to centrifugation at 20,000g, 4°C for 90 min. Soluble fractions were collected and stored at -80°C for downstream MS processing in TMT-16plex experiments.

### PISA Mass Spectrometry Sample Processing

15 μg of soluble protein fraction was removed after centrifugation and placed in 200μL PCR tubes. Dilution buffer (400mM HEPES pH 8.5, 2% SDS) was added to each sample (protein denaturation). TCEP was added to 5mM and incubated for 30 minutes (protein reduction). Iodoacetamide was added to 10mM and incubated at room temperature for 30 minutes (alkylation). DTT was added to 10mM (quench) and incubated for 15 minutes. Cleanup was performed using SP3 beads (Cytiva, 1:1 E3:E7 mix) using 10 μg of beads per μg protein. 3 μL of beads at 50 μg/μL were added to each sample and mixed by pipetting. Ethanol was added to final 50% and incubated with shaking for 15 minutes. PCR tubes were placed on magnetic rack and unbound supernatant was discarded. Beads were washed three times with 80% Ethanol. Final wash supernatant was removed and beads were air dried off the magnet. 100 μL of digestion solution (50mM TEAB pH 8.0 containing 150 ng Trypsin/LysC mix (Promega V5073)) was added to each sample and incubated at 37°C overnight with shaking at 1000 rpm. After digestion, PCR tubes were placed on a magnet and the supernatant was removed. Samples were TMT-labeled (Thermo A44521) using 2:1 TMT label:protein ratio for 45 min at RT. TMT labeling reactions were quenching with hydroxylamine prior to combining 16 reactions. Samples were de-salted and fractionated using High pH Reversed-Phase Peptide Fractionation Kit (Thermo 84868) prior to speedvac and LC-MS/MS analysis.

### LC-MS/MS operation

16plex-TMT samples for PISA, TRAP samples, and TSC22D4 UV-C samples were run on Oribtrap Eclipse in 180 minute gradient with parameters as follows. In MS1, scan range of 400-1600 m/z, maximum injection time of 50 ms, AGC target of 4e5, mass tolerance 10ppm. In MS2, CID collision energy of 35%, AGC target 1e4, mass tolerance 25 ppm. For MS3, isolation window 0.7, collision energy 35%, orbitrap resolution 50K, maximum injection 200ms, AGC target 1e5. IGF2BP3-UVC samples were run in a 190 minute method on a Fusion Lumos (Thermo). For MS1, parameters were as follows: 60K Orbitrap resolution, 300-1200m/z scan range, 50ms max injection time, AGC target 4e5. For MS2, HCD with collision energy 30%, max injection time 35 ms, AGC target 1e4.

### Thermal Shift Western

Lenti-X 293T cells (Takara 632180) were seeded into 15-cm plates at 70% confluency. For IGF2BP2 and TSC22D4, cells were transfected the next day with 20 μg of DNA (encoding expression vector for IGF2BP2 or TSC22D4) per plate and 6.6 μg PEI MAX (Polysciences). Cells were harvested in PISA lysis buffer. Lysate was diluted to 2.2mg/mL and split across 16 tubes and treated with 2 mM glucose or vehicle. Samples briefly spun down in microcentrifuge and incubated at room temperature for 30 minutes without rotation or shaking, followed by separating each sample into 8 tubes for heating in thermal cycler at 40, 44, 48, 52, 56, 60, or 64°C for 3 min before moving to ice bucket. Samples were lysed by adding 4X LDS + BME and run on western blot.

### Recombinant protein production

Lenti-X 293T cells (Takara 632180) were seeded into 15-cm plates at 70% confluency. The next day, cells were transfected with 20 μg of DNA per plate and 6.6 μg PEI MAX (Polysciences). Cells were harvested 48-72 hours post transfection. Cells were lysed in Flag lysis buffer (50mM Tris-HCl pH 7.5 150mM NaCl, 1mM EDTA, 1% Triton-X, and 1X Protease Inhibitor Cocktail [Sigma, P8340]) 30 minutes on ice, then clarified by centrifugation at 20,000xg. Lysate was incubated with anti-FLAG M2 affinity gel (MilliporeSigma A2220) 2 hours to overnight, then washed twice with Flag wash buffer (50mM TrisHCl pH 7.5, 3 mM EDTA, 0.5%NP-40, 500 mM NaCl, 10% Glycerol) and once with PBS before elution in PBS with 0.5 mg/mL Flag peptide. Eluates were dialysed in PBS overnight using Slide-A-Lyzer™ MINI Dialysis Device, 7K MWCO (Thermo 69562). Optionally, purified protein was concentrated in 10K MWCO Amicon columns (Millipore UFC500396). Finally, the target protein concentration was determined by running purified protein and a BSA standard curve on a 10% Bis-Tris gel and staining using InstantBlue Coomassie Protein Stain (Abcam ab119211).

### Microscale Thermophoresis (MST)

Recombinant protein was labeled at 100nM using Monolith His-Tag labeling Kit RED-tris-NTA (Nanotemper) using 1:2 protein:dye ratio. Prior to labeling protein was quantified by BSA standard curve in SDS-PAGE, and if necessary, protein was buffer exchanged to remove any primary amine buffer components. A 16-part dilution series (2x dilutions) of the ligand was made and labeled protein was mixed 1:1 with the ligand and incubated for 5 minutes at room temperature prior to loading into Monolith NT.115 Capillaries or NT.115 Premium Capillaries. MST was performed using a Monolith NT.115 instrument (NanoTemper) at room temperature, using 60% excitation power and medium MST power.

### UV-C Mass Spectrometry

Glucose UV-C crosslinking mass spectrometry was performed as previously described.^4^ Briefly, recombinant protein was diluted to 200 nM and incubated with glucose (10 mM for TSC22D4, 20 uM for IGF2BP3) at room temperature for 15 minutes, aliquoted into 3 uL droplets on parafilm, and subjected to 254 nm UV-C light (UV Stratalinker 2400) at 0.3 J/cm^2^. All droplets for each sample were combined, boiled in LDS, and subjected to in-gel digestion prior to LC-MS/MS analysis.

### Target Response Accessibility Profiling (TRAP)

3 μg of Recombinant TSC22D4 was incubated with 10mM glucose or vehicle for 30 minutes at room temperature. Deuterium Formaldehyde (CD20) was added to a final concentration of 0.03% and Borane Pyridine Complex (BPC) was added to 3.65mM. Reaction was incubated for 30 minutes at room temperature prior to quenching with 50mM NH4CO3. Protein was processed for mass spectrometry using Filter-Aided Sample Preparation (FASP) method. Briefly, sample was filtered using 10 kDa column at 12000g spin for 15 minutes. Protein was denatured using 8M Urea for 30 min, reduced with 5mM DTT at 56°Cfor 30 minutes, and alkylated with 20mM IAA for 30 min at room temperature. 10 mM DTT was added to quench IAA and protein was washed twice with NH4CO3. 60 ng trypsin per sample was added in 50mM TEAB digestion buffer. Peptides were collected from the column and de-salted with monospin C18 prior to LC-MS/MS analysis.

### BS3 Crosslinking Mass Spectrometry

2.5 μg of recombinant TSC22D4 protein was crosslinked at 2uM protein concentration using 50 μM BS3 in the presence of 10mM glucose or vehicle (PBS) at room temperature for one hour prior to quenching with 4X LDS sample buffer (ThermoFisher NP0008). The monomeric band was cut from the gel and processed by in-gel digestion. Thermo .raw files were converted to .mgf files prior to search with XiSearch (v 1.7.6.1) and filtered by XiFDR (v. 2.1.5.2)^46^.

### In-gel digestion

Protein samples were boiled with 4X LDS sample buffer (ThermoFisher NP0008), run in a 10% Bis-Tris gel, and stained with InstantBlue (Abcam ab119211). The protein band of interest was cut from the gel and diced into roughly 1 mm squares. InstantBlue stain was removed with overnight incubation in 50% ACN. Samples were then reduced using 20mM DTT in 50mM ammonium bicarbonate (ABC) at 37°C for one hour. Samples were alkylated using 55mM IAA in 50mM ABC, room temperature in the dark for 30 minutes. Gel slices were washed three times with 50mM ABC. Gel slices were then washed with 100% Acetonitrile and air dried. Protein was digested overnight at 37°C using 1:50 reagent:protein Trypsin/Lys-C in 50mM ABC and 0.02% Protease Max surfactant (Promega). Peptide was extracted from digest liquid, two elutions with 5% acetic acid, and two elutions with 2.5% acetic acid, 50% acetonitrile. Elutions were pooled and any extra SDS-Page gel was removed using Spin-X 0.22uM column prior to speedvac and de-salt using MonoSpin columns (GL Sciences) and analysis by mass spec.

### Lentivirus Production, Infection, and Selection

Approximately 6 million Lenti-X™ 293T (Takara Bio #632180) were plated in 10-cm dish, then transfected the next day with 7.5 μg pLEX or pLKO.1 expression vector, 7.5 μg p8.91 (Addgene #187441), and 3.75 μg pMDG (Addgene #187440) combined with 50 μg polyethylenimine in 2 mL OptiMEM (Thermo #11058021). After 2 days, supernatant was collected and filtered through 0.45 micron filter, then aliquoted and stored at -80°C. For infections of beta cells, lentivirus was concentrated with Lenti-X Concentrator (Takara #631231) according to manufacturer directions before freezing. Cells were infected with thawed lentivirus and 5 ug hexadimethrine bromide (Sigma H9268) per mL media. Media was changed after 1 day and cells were selected 2-3 days after infection. 3T3-L1 were selected with 3 ug/mL puromycin for 2 days. MIN6 and MIN6-6 cells were selected with 1 ug/mL puromycin for 3 days or 1 ug/mL blasticidin for 5 days. Lenti-X 293T cells were selected with 2 ug/mL puromycin for 3 days.

### Fast-Protein Liquid Chromatography (FPLC)

FPLC was performed as previously described.^4^ Lenti-X 293T cells stably infected pLEX FHH-TSC22D4 were incubated with glucose-free DMEM (Thermo 11966025) supplemented with 10% FBS and 1% PenStrep overnight. Cells were collected via trypsinization and washed twice in phosphate-buffered saline (PBS). The pellet was lysed in 10 pellet volumes of nuclei lysis buffer (20 mM HEPES, pH 7.5, 300 mM KCl, 5 mM MgCl2, 0.5 mM DTT, 1X PIC) and incubated on ice for 5 minutes. Lysates were split into two samples, and glucose was added to achieve the desired final concentration. Benzonase (Sigma E1014) was then added to reach a concentration of 3-5 Units/µL, and the samples were rotated at room temperature for 30 minutes. The samples were centrifuged at maximum speed at 4°C for 10 minutes. The resulting supernatant (250 µL) was injected a 200 uL sample loop on AKTA machine equipped with Superose 6 Increase 10/300 column (Cytiva 29-0915-96, 10/300 GL) and Fraction collector F9-C, and the specified program was initiated. A mixture of thyroglobulin, g-globulin, ovalbumin, myoglobin, and vitamin B12 was used as gel filtration standards (BioRad, #1511901).

### Multiple sequence alignment

UniProt canonical amino acid sequences of TSC22D1, TSC22D2, TSC22D3, and TSC22D4 were inputted into Clustal Omega^39^ and visualized using Jalview^40^.

### CUT&RUN

CUT&RUN was performed on 3T3-L1 cells virally infected with HA-tagged TSC22D4 using reagents from the EpiCypher kit (EpiCypher, 14-1048). 3T3-L1 cells were washed with PBS, harvested with trypsin, quenched with DMEM, and resuspended in cold PBS + 0.1% BSA to be maintained afterwards at 4°C. Nuclei were extracted by resuspending in 100 μL nuclei extraction buffer per 1-2.5 M cells (20 mM HEPES pH 7.5, 10 mM NaCl, 0.5 mM spermidine, 0.1% BSA, 0.1% NP-40, and 1 tablet Roche mini protease inhibitor per 10-15 mL) and incubating for 10 minutes on ice. 5-10 volumes of wash buffer (20 mM HEPES pH 7.5, 150 mM NaCl, 0.5 mM spermidine, 0.1% BSA, 0.05% Triton X-100, and Roche protease inhibitor) were added, cells centrifuged at 600 RCF, and supernatant removed. After an additional wash, cells were quantified using a Countess. 500K nuclei were bound to 10 μL activated Concanavalin A beads (EpiCypher, 21-1401) per sample for 10 min incubation at 4°C. After removing supernatant, beads were incubated with rocking at 4°C for 2 hrs with 1 µg antibody [either anti-HA antibody (Cell Signaling, 3724S) or IgG control antibody (Cell Signaling, 3900S)] diluted in 50 μL Antibody Buffer (wash buffer with 2 mM EDTA). After staining, cells were washed twice with wash buffer, and incubated with 1.7 μL pAG-MNase (EpiCypher, 15-1016) per sample for 1 hour at 4°C with rocking, then washed with once with 150 μL wash buffer, then 150 μL low salt buffer (20 mM HEPES pH 7.5, 0.5 mM spermidine, 0.1% BSA, 0.05% Triton X-100, and Roche protease inhibitor) and resuspend in 50 µL low salt buffer. After 5 minutes on ice, 2 μL of 250 mM CaCl2 was added to the beads and beads were subsequentially incubated on ice for 20 minutes. The reactions were quenched with 17 μL 4X STOP buffer (600 mM NaCl, 40 mM EDTA, 80 mM EGTA, 0.05% SDS, 1% Triton X100, 0.1 mg/mL RNAse A, 0.1 mg/mL glycogen, 0.6 pg/uL spike-in E. coli DNA) for 30 minutes at 37°C according to the manufacturer’s protocol. Beads were placed on a magnet, and supernatant containing released fragments was removed. DNA was extracted by a Zymo DNA Clean and Concentrator (Zymo Research, D4014), eluting in 27 μL water. Library was prepared using the NEB Ultra II DNA library prep kit (NEB, E7103S) following published protocol^47^, with the following modifications: the adapter was diluted 20-fold, PCR was performed with SYBR dye to visualize amplification, and after PCR, 20 μL of water, then 30 μL of beads (0.6X instead of 0.8X) beads were added for the initial selection to remove large small fragments, followed by the second size selection using the addition of 20 μL beads (1X instead of 1.2X) for library capture. Sequencing was performed on a NovaSeq X Plus, paired-end, 150 bp length, with at least 20 M reads for each sample.

### ChIP

Chromatin immunoprecipitation (ChIP) assays were performed as previously described with minor modifications.^48^ Briefly, 3T3-L1 cells were cross-linked with 1% formaldehyde (Promega) and cells were lysed in nuclear lysis buffer [0.1M Tris, pH8, 1% SDS, 10mM EDTA]. Chromatin was sonicated to an average fragment length of 150-250 bp using a Bioruptor (Diagenode). Samples were diluted in ChIP dilution buffer [0.01% SDS, 2% Triton X-100, 16.7mM Tris pH 8, 167mM NaCl], pre-cleared with Protein G Dynabead (ThermoFisher), incubated with IgG or HA antibody (Cell Signaling) and chromatin was immunoprecipitated overnight at 4°C. Antibody capture was performed with Protein G dynabeads for 2 hours. Samples were then washed three times in ChIP wash buffer. Following cross-link reversal DNA was purified using ChIP DNA Clean and Concentrator Kit (Zymo). Purified ChIP DNA was quantified using the high sensitivity DNA assay and Qubit fluorometer (Thermo).

ChIP-qPCR was performed with Lcn13 primers previously described.^10^ Mouse Negative Control Primer Set 1 (Active Motif #71011) was used as control.

### Affinity Purification Mass Spectrometry

3T3-L1s stably expressing HA-tagged TSC22D4 were grown to 90% confluence, washed once with PBS, then treated with glucose-free DMEM (Thermo 11966025) + 10% FBS for 4 hours, incubated 1 hour with DMEM 10% FBS with 25 mM glucose or 25 mM 3-OMG, then collected by trypsinization. Cell pellets were stored at -80C. Pellets were lysed on ice for 30 minutes in 5X pellet volume with 50 mM HEPES, pH 7.5, 100 mM NaCl, 0.2 mM MgSO4, 1% Triton X-100, 1X Halt Protease/Phosphatase Inhibitor (Thermo 78440), and clarified by centrifugation (20,000xg for 10 min at 4°C). Supernatant was collected and quantified by BCA assay. Added BS3 (Thermo A39266) to final 1 mM concentration and let sit 30 minutes at RT. 0.4 mg of lysate was used for IP, and IP volume made up to 1 mL lysis buffer. 20 uL of Pierce HA Dynabeads were added and rotated at 4C overnight. Beads were washed three times with 500 uL of IP buffer for 5 minutes each. Proteins were eluted off beads in 30 uL 2X LDS sample buffer with 90 degC heating for 5 min. Samples were then run on SDS page and processed with in-gel digestion for mass spec.

MaxQuant^49^ (v. 2.4.7.0) was used to process LC-MS/MS data for protein identification and quantification. Spectra were searched against the reviewed *Mus musculus* Uniprot database, with decoy sequences generated in revert mode and common contaminants included. Carbamidomethylation of cysteine was set as a fixed modification, while oxidation of methionine and N-terminal acetylation were used as variable modifications. A minimum peptide length of 7 amino acids was required. Mass tolerance for both MS and MS/MS was set to 20 ppm for FTMS and 0.5 Da for ITMS. False discovery rates were controlled at 1% for PSMs, proteins, and modification sites.

### Gene Ontology for AP-MS

Using geneontology.org, with SAINT score threshold of 0.7 and reference list of all detected proteins in mass spec.

### StringDB

Genes were considered condition-specific if SAINT score above 0.7 and FC was at least 2X other condition. For glucose-enriched proteins, both glucose and 3MG conditions were required to be 2-fold enriched over vehicle.

### Mitochondrial Purification

Mitochondria were isolated as previously performed with modifications.^50^ 15-cm plates of 3T3-L1s grown to ∼80% confluence, treated 4 h with glucose-free DMEM + FBS then either collected (starve) or incubated 1 h with standard DMEM + FBS (25 mM glucose). 2 15-cm plates of 3T3-L1 collected by scraping in total 2 mL mitochondrial isolation buffer (20 mM HEPES, 220 mM mannitol, 70 mM sucrose, 1 mM EDTA, 1X Protease Inhibitor Cocktail), incubated on ice for 20 min and homogenized manually. Starved cells lysed much more easily and 100 strokes yielded >90% Trypan blue positive cells. Glucose treated cells required 150 strokes for ∼80% Trypan blue positive. Supernatant clarified at 800 g for 10 min was pelleted at 10,000 g for 20 min yielding a crude mitochondrial fraction. The pellet was resuspended in mitochondrial isolation buffer, clarified at 800 g for 20 min and pelleted at 10,000 g for 20 min resulting in an enriched mitochondrial fraction. Mitochondrial fraction lysed in RIPA.

### Protein-Metabolite Co-Folding

Boltz-2^32,51^ was run on Rowan Labs (https://labs.rowansci.com/) using AlphaFold structure^36,52,53^ for IGF2BP2 or IGF2BP3 and beta-D-glucose.

### IGF2BP3 easyCLIP

easyCLIP was performed and analyzed as published^54^ using Lenti-X 293T expressing HA-IGF2BP3.

### Glucose Stimulated Insulin Secretion (GSIS) Assay

100,000 MIN6 or MIN6-6 cells were seeded per 48-well. The next day, cells were incubated 2h with glucose-free Krebs-Ringer Bicarbonate Buffer (KRBH) then 1h with either 200 μL KRBH supplemented with 2.8 mM glucose or 16.7 mM glucose. Conditioned media was collected and debris and cells were removed by spinning 500xg for 3 min. For MIN6 cells, insulin content was collected by rocking at 4C overnight in 100 μL acid ethanol and quenched with 100 μL 1M Tris pH 7.5. Supernatant and insulin content samples were diluted (1:7 for low glucose, 1:20 for high glucose, and 1:500 for content) and 10 μL was input to Mercodia Mouse Insulin ELISA (Mercodia 10-1247) and run according to manufacturer instructions. For MIN6-6 cells, 10 μL supernatant was used in luciferase assay using Promega Nano-Glo® Luciferase Assay System (Promega N1110) according to manufacturer instructions.

### RIP-qPCR and RIP-seq

MIN6-6 cells or HIT-T15 pancreatic beta cells were starved in KRBH (5 mM KCl, 120mM NaCl, 15mM HEPES pH 7.5, 25 mM NaHCO_3_, 1mM MgCl_2_, 2mM CaCl2, 1 mg/mL BSA) for 2 hours, then treated with 16.7 mM Glucose or PBS for 1.5 hours. Cells were washed with PBS and lysed on plate with RIP lysis buffer (140 mM KCl, 1.5 mM MgCl2, 20 mM Tris-HCl pH 7.4, 0.5% NP-40, 0.5 mM DTT, 1 U/mL RNase inhibitor, protease inhibitor), lysate was cleared at 15000g for 10 min at 4C. Lysates were precleared with 20μL Protein G magnetic beads (Pierce) for 30 min at 4°C with end over end rotation. 25 μL of Protein G beads pre-coated with 1.5μg of IMP2 antibody (ProteinTech) or HA antibody (Cell Signaling) were added to the lysates and incubated for at least 2 hours at 4°C with end over end rotation. Beads were washed four times with lysis buffer prior to elution with RLT + BME. RNA was collected with RNeasy mini column for RT-qPCR and protein flow through was precipitated with acetone for western blot and mass spec analysis. qPCR was performed as described above, with half of the immunoprecipitaiton used as input for the cDNA synthesis. RIP-seq libraries were prepared from RNA using Takara SMARTer® Stranded Total RNA-Seq Kit v3 - Pico Input Mammalian (cat #634489) according to the manufacturer’s protocol. RIP-seq libraries were sequenced on the Illumina NovaSeq X Plus platform, with at least 20 M paired-end 150 bp reads per sample.

### IGF2BP2 Conservation

Orthologous IGF2BP2 protein sequences were assembled by querying the UniProt API^55^ using the reviewed human IGF2BP2 entry (UniProt: Q9Y6M1) as the reference and searching species records via gene- and protein name–based strategies, followed by retrieval of FASTA sequences for the top-matching accessions. These sequences were subjected to multiple sequence alignment using MAFFT^56^ in automatic mode to infer homologous positional correspondence across taxa. The resulting aligned FASTA was parsed for downstream per-site analyses, and alignment coordinates were mapped to ungapped query (human) coordinates by tracking non-gap positions in the query sequence, enabling interpretation of alignment-derived site wise statistics in the reference protein’s numbering.

For each alignment column, we computed (i) an “identity” metric defined as the frequency of the most common non-gap residue (i.e., consensus frequency), (ii) Shannon entropy^57^ over the non-gap residue distribution as a measure of positional variability, and (iii) the fraction of sequences containing a gap at that site. Entropy is further normalized by the theoretical maximum for a 20-letter amino-acid alphabet (log220) to yield a scaled variability index, and a conservation score is defined as 1 - Entropynorm, so that higher values indicate stronger conservation.

Region-focused summaries are then computed for predefined query-coordinate intervals (3–7, 40–44, 70–77), reporting mean identity, mean conservation score, mean entropy, and counts of highly conserved sites (>0.9 identity) and variable sites (<0.5 identity).

---

**Figure S1.**
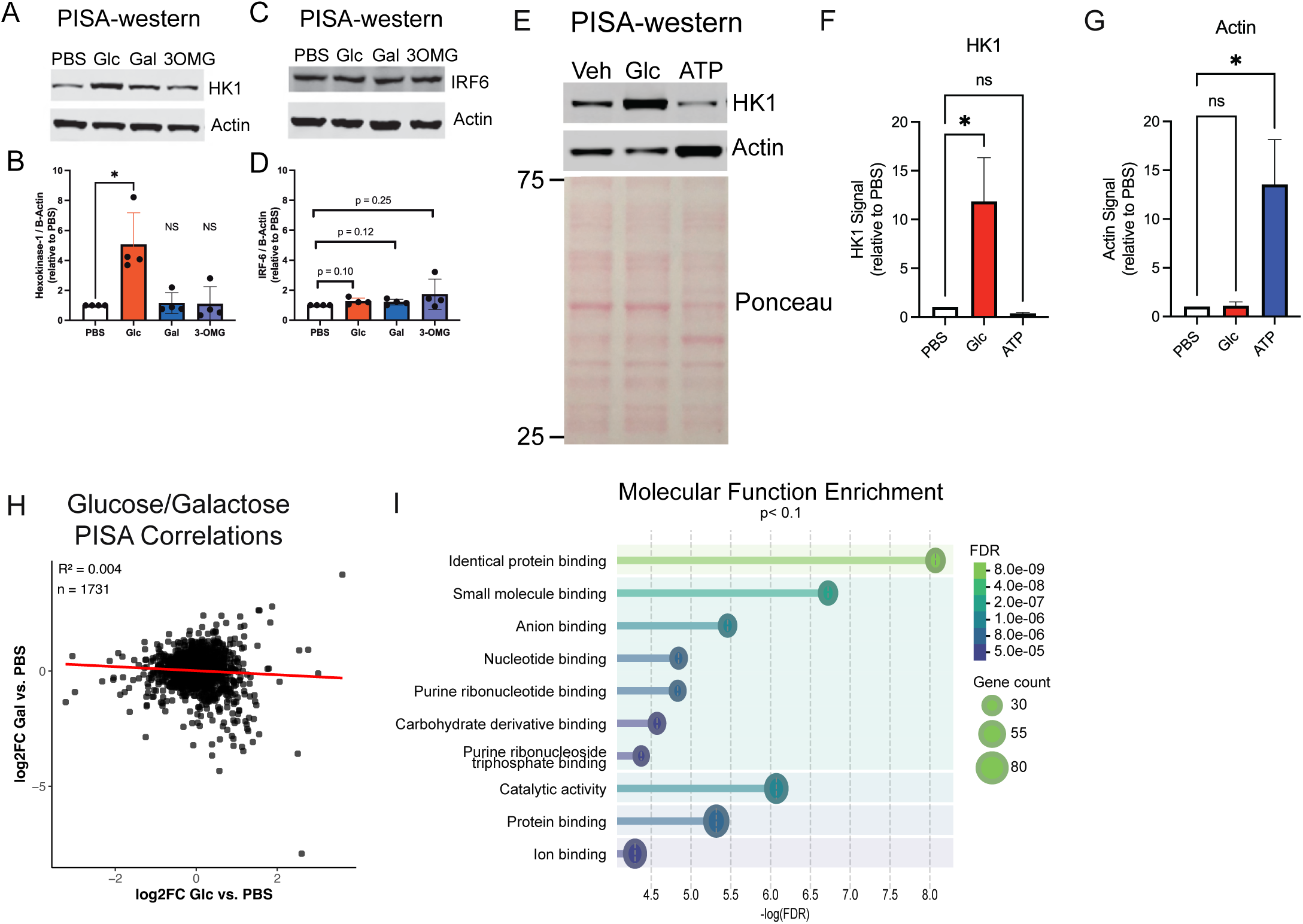
Thermal profiling proteomics identifies glucose binding proteins at physiologic glucose concentrations. **A**) Western blot showing solubilization of glucose-binding protein HK1 with glucose but not galactose or 3MG, quantified in **B. C**) Western blot showing lack of solubilization of glucose-binding protein IRF6, quantified in **D**. **E**) Western blot and Ponceau S stain comparing solubilization of glucose-binding protein HK1 with glucose and ATP-binding protein actin with ATP and total proteome solubilization, quantified in **F** and **G**. **H**) Correlations between glucose and galactose treatments on thermal solubility in PISA assay in 3T3-L1 pre-adipocyte lysates. **I**) Gene ontology analysis of proteins nominated by PISA with p<0.10.

**Figure S2.**
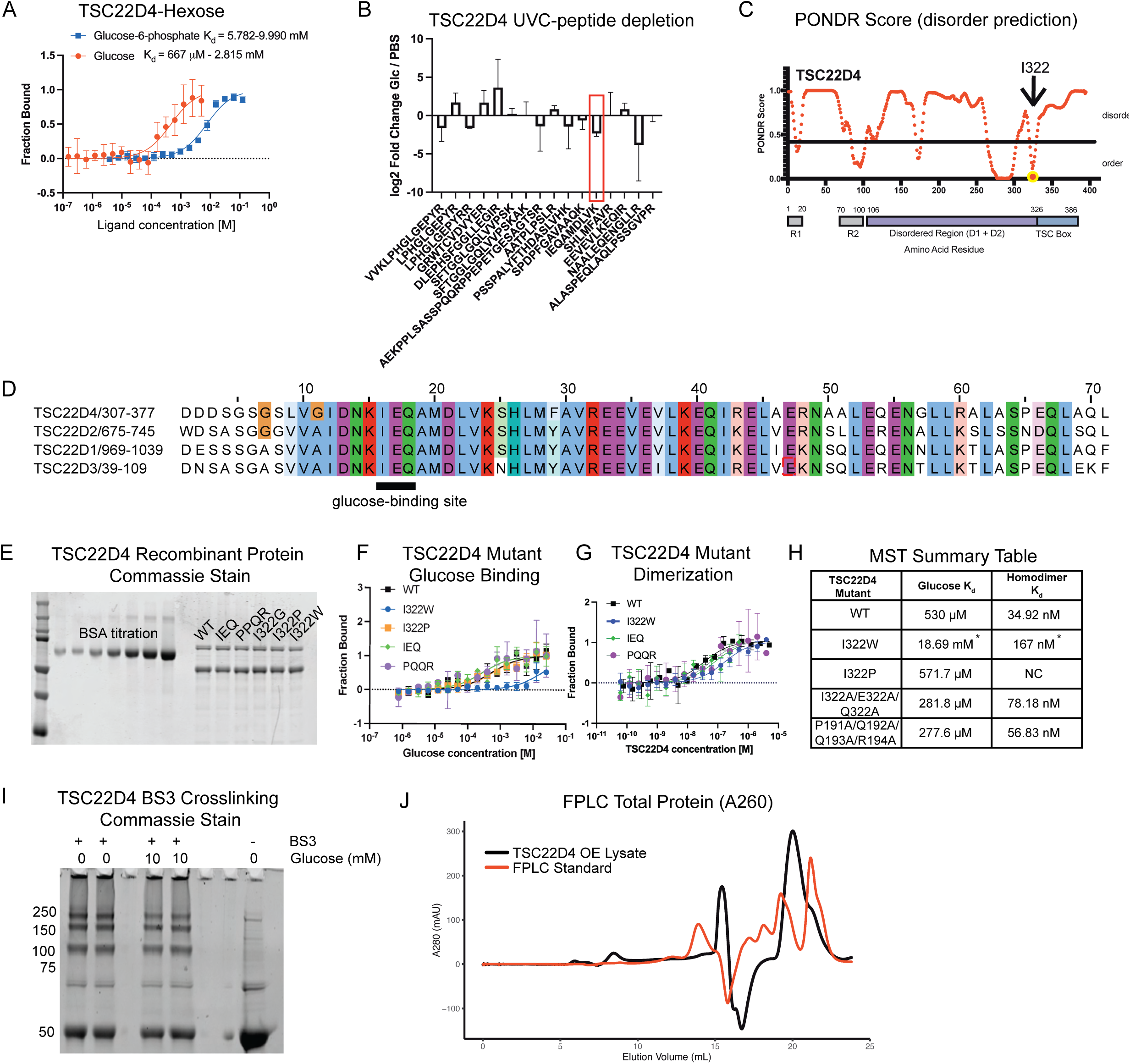
TSC22D4 binds glucose at TSC box to increase C-terminal accessibility. **A)** MST analysis of TSC22D4 affinity toward glucose vs. glucose-6-phosphate. **B**) Glucose-induced changes in unmodified peptide abundance from UVC-MS experiment shows loss of unmodified IEQAMDLVK peptide (boxed in red). Glucose binding events on peptides are often seen with loss of unmodified counterparts. **C**) PONDR predicts I322 occurs at an ordered protein region directly upstream of the TSC box. Domains are annotated based on domains proposed in Demir et al 2022 and mapped to human sequence. **D**) Multiple sequence alignment of human TSC22 domain family proteins using Clustal Omega^39^ and visualized using Jalview^40^. **E**) Coomassie blue SDS-page analysis of purified TSC22D4 protein variants. **F**) Glucose binding activity of TSC22D4 mutant panel. **G**) Dimerization of TSC22D4 mutant panel **H**) Calculated dissociation constant from MST in **F** and **G**, with * representing values significantly different from WT (p<0.05 using Extra sum-of-squares F Test). **I**) SDS-PAGE showing BS3 crosslinking in TSC22D4. Monomeric band isolated for mass spec runs around 50 kDa. **J**) A260 readout of TSC22D4 FPLC experiment compared with FPLC standard.

**Figure S3.**
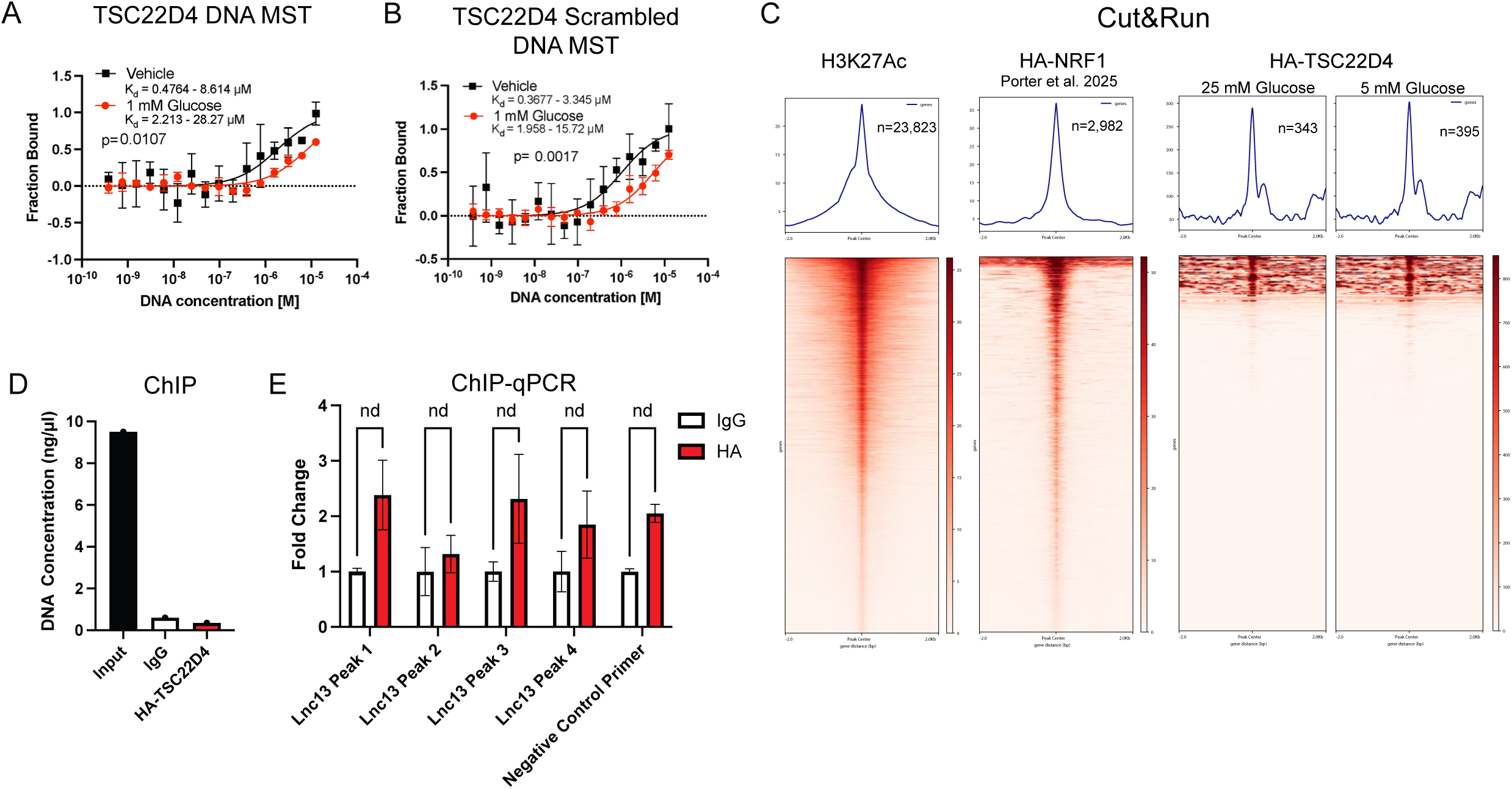
No detectable TSC22D4-chromatin interaction identified in 3T3-L1 cells by Cut&Run and ChIP. **A**) MST analysis of TSC22D4 with DNA oligo synthesized to match genomic region containing TSC22D4 binding motif obtained Homer^41^ motif analysis of Encode accession ENCSR787RVK^42–44^. **B**) MST analysis of TSC22D4 with scramble DNA oligo. **C**) Cut&Run peak-centered heatmap of H3K27Ac and HA-TSC22D4 pulldown, showing technical success of Cut&Run experiment, compared with previously generated Cut&Run data of NRF1 using identical antibody and protocol as TSC22D4^45^. **D**) DNA quantification from HA-TSC22D4 ChIP in 3T3-L1. **E**) ChIP-qPCR of HA-TSC22D4 in 3T3-L1 using primers for genomic regions bound by TSC22D4 in HepG2^10^

**Figure S4.**
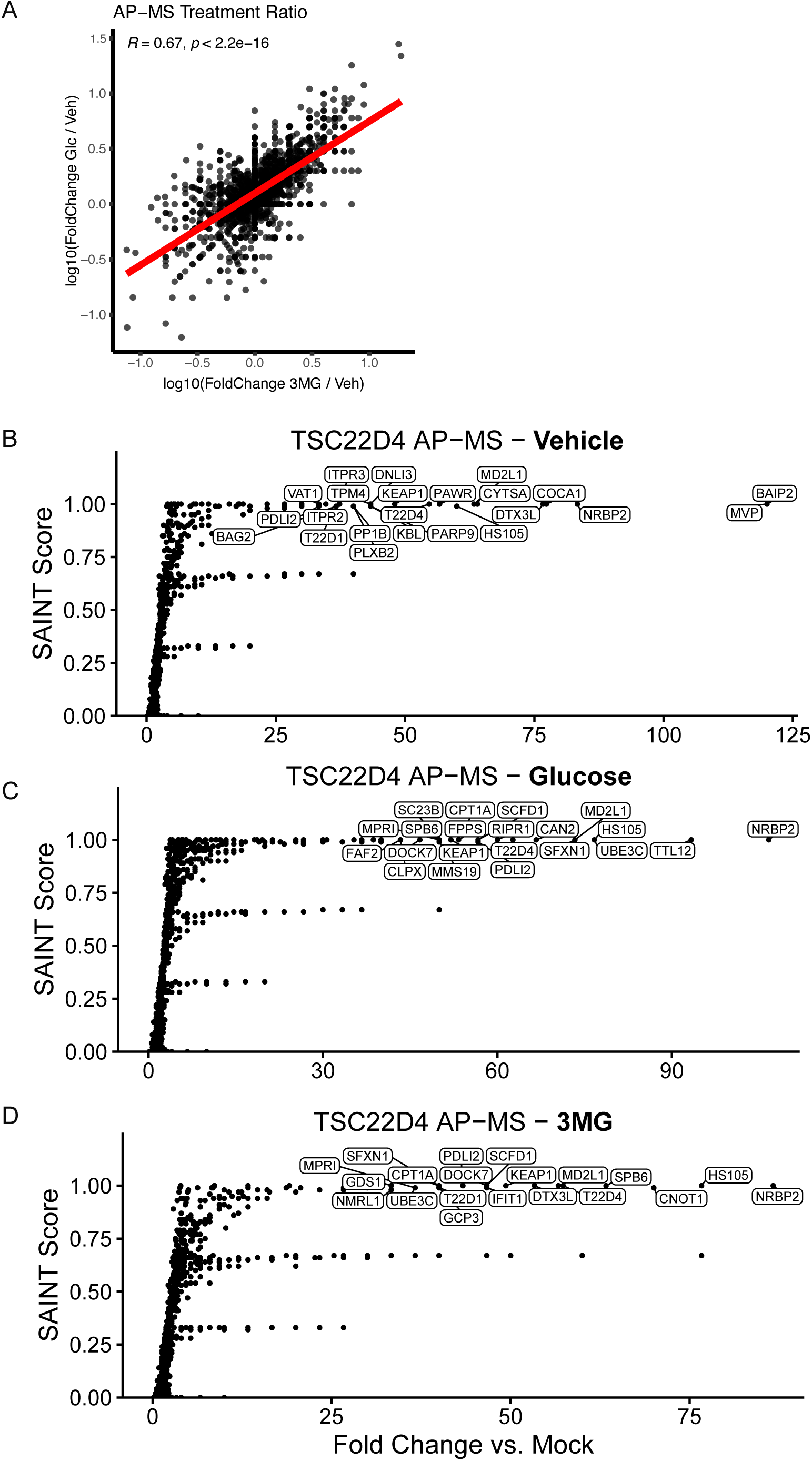
Glucose modulates TSC22D4 interactome. **A**) Correlation between enrichment of prey proteins in glucose and 3MG treatment (relative to vehicle) in TSC22D4 AP-MS **B,C,D**) TSC22D4 AP-MS data performed in 3T3-L1 in vehicle (**B**), glucose (**C**), or 3MG (**D**) conditions.

**Figure S5.**
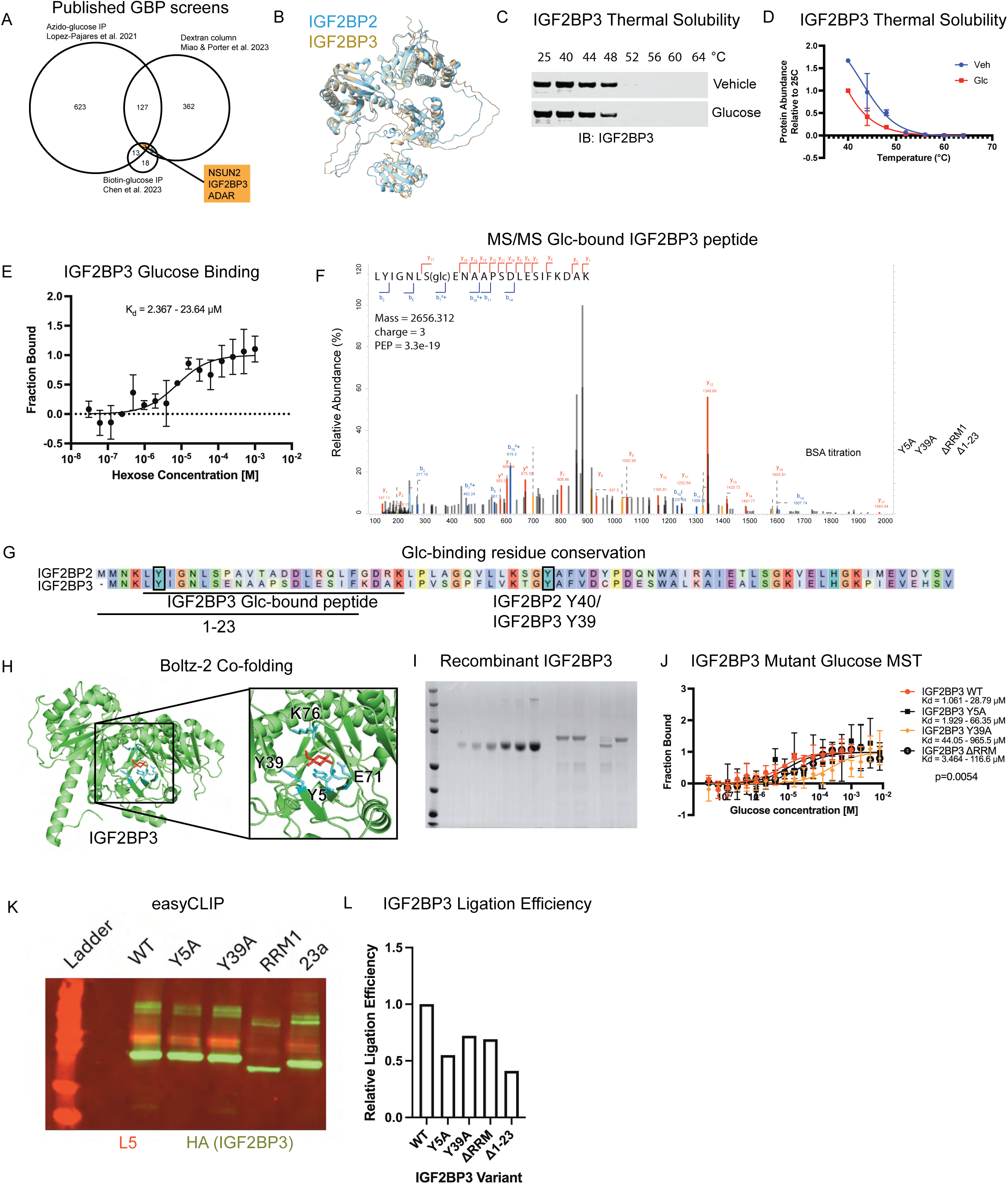
IGF2BP3 binds glucose at RRM1 domain. **A**) Intersection of glucose binding protein mass spec experiments showing NSUN2, IGF2BP3, and ADAR as nominated glucose binding proteins. **B**) Overlay of AlphaFold models of IGF2BP2 and IGF2BP3 proteins^36–38^. **C**) Thermal solubility assay western profile for IGF2BP3 in HEK293T. **D**) Quantification of thermal solubility assay in C. **E)** IGF2BP3-Glucose MST analysis showing similar affinity for glucose as IGF2BP2. **F**) MS/MS spectrum of glucose-linked peptide at N-terminal region of IGF2BP3. **G**) Sequence alignment of N-terminus of IGF2BP2 and IGF2BP2 annotated with glucose-bound peptide, N-terminal amino acids deleted in mutant panel, and IGF2BP2 Y40 (synonymous with IGF2BP3 Y39) **H**) Boltz-2 co-folding of IGF2BP3 and glucose with residues contacting glucose highlighted in cyan **I**) Coomassie gel showing recombinant protein variants of IGF2BP3. **J**) MST analysis of IGF2BP3 mutant panel showing Y39A mutation ablates glucose binding **K**) easyCLIP ligation efficiency analysis of IGF2BP3 variants. **L**) Quantification of easyCLIP ligation efficiency

**Figure S6.**
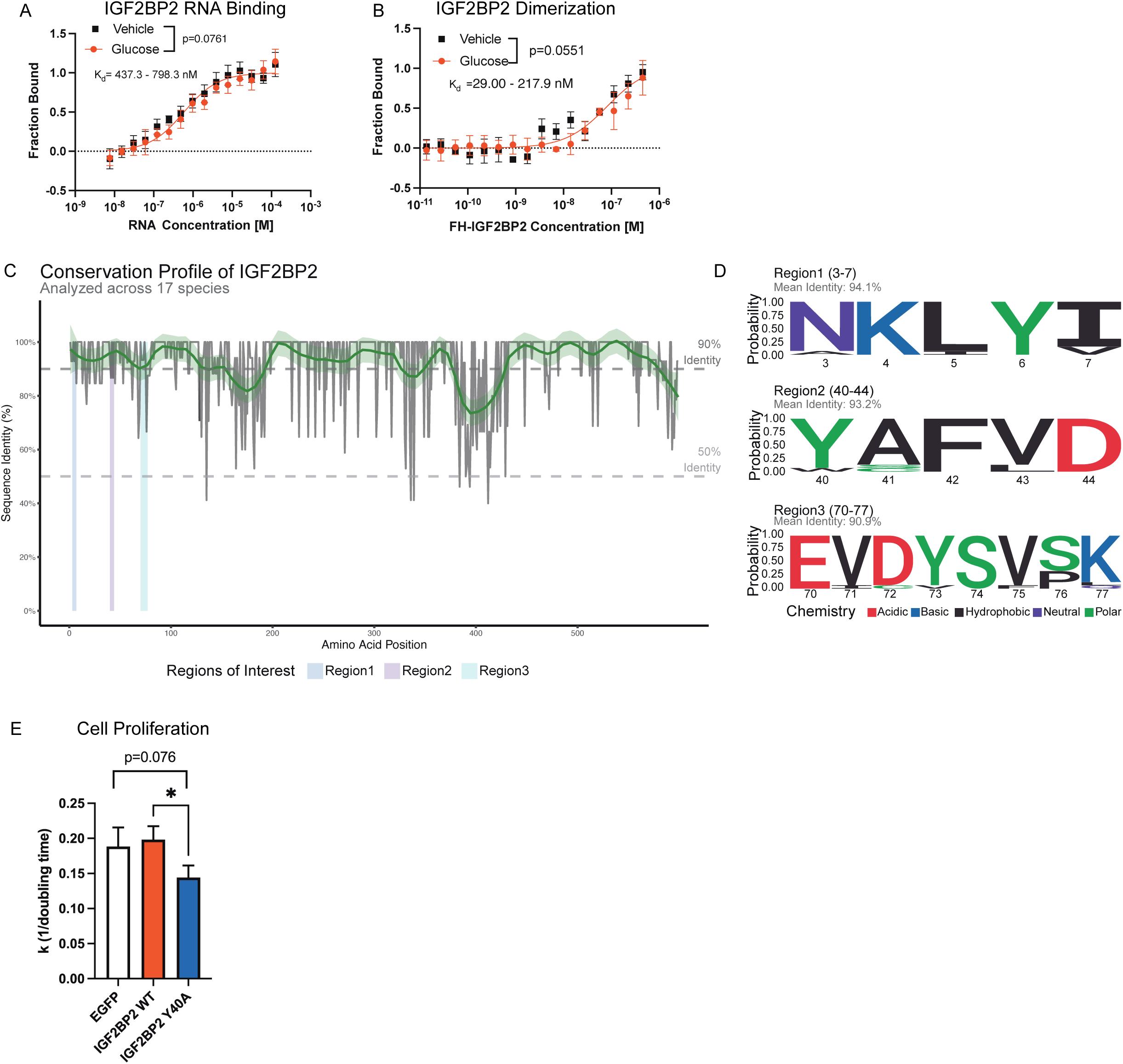
IGF2BP2 binds glucose at RRM1 domain to modulate insulin secretion. **A**) MST analysis of IGF2BP2 with RNA oligo shows no significant difference in RNA binding upon glucose treatment **B**) MST analysis shows no significant difference in homodimerization upon glucose treatment **C**) Conservation analysis of IGF2BP2 orthologs shows glucose-binding regions are well-conserved across 150 million years of evolution **D**) Sequence homology among IGF2BP2 orthologs at glucose-binding regions **E**) Cell-titer-blue proliferation assay in MIN6-6 cells showing decrease in proliferation rate in Y40A overexpression cells.

## References

1. Liu, W., Wang, Y., Bozi, L.H.M., Fischer, P.D., Jedrychowski, M.P., Xiao, H., Wu, T., Darabedian, N., He, X., Mills, E.L., et al. (2023). Lactate regulates cell cycle by remodelling the anaphase promoting complex. Nature 616, 790–797. 10.1038/s41586-023-05939-3.

2. Mitro, N., Mak, P.A., Vargas, L., Godio, C., Hampton, E., Molteni, V., Kreusch, A., and Saez, E. (2007). The nuclear receptor LXR is a glucose sensor. Nature 445, 219–223. 10.1038/nature05449.

3. Lopez-Pajares, V., Bhaduri, A., Zhao, Y., Gowrishankar, G., Donohue, L.K.H., Guo, M.G., Siprashvili, Z., Miao, W., Nguyen, D.T., Yang, X., et al. (2025). Glucose modulates IRF6 transcription factor dimerization to enable epidermal differentiation. Cell Stem Cell 32, 795–810.e10. 10.1016/j.stem.2025.02.017.

4. Miao, W., Porter, D.F., Lopez-Pajares, V., Siprashvili, Z., Meyers, R.M., Bai, Y., Nguyen, D.T., Ko, L.A., Zarnegar, B.J., Ferguson, I.D., et al. (2023). Glucose dissociates DDX21 dimers to regulate mRNA splicing and tissue differentiation. Cell 186, 80–97.e26. 10.1016/j.cell.2022.12.004.

5. Chen, T., Xu, Z.-G., Luo, J., Manne, R.K., Wang, Z., Hsu, C.-C., Pan, B.-S., Cai, Z., Tsai, P.-J., Tsai, Y.-S., et al. (2023). NSUN2 is a glucose sensor suppressing cGAS/STING to maintain tumorigenesis and immunotherapy resistance. Cell Metab. 35, 1782–1798.e8. 10.1016/j.cmet.2023.07.009.

6. Miao, W., Porter, D.F., Li, Y., Meservey, L.M., Yang, Y.-Y., Ma, C., Ferguson, I.D., Tien, V.B., Jack, T.M., Ducoli, L., et al. (2024). Glucose binds and activates NSUN2 to promote translation and epidermal differentiation. Nucleic Acids Res. 52, 13577–13593. 10.1093/nar/gkae1097.

7. Gaetani, M., Sabatier, P., Saei, A.A., Beusch, C.M., Yang, Z., Lundström, S.L., and Zubarev, R.A. (2019). Proteome Integral Solubility Alteration: A High-Throughput Proteomics Assay for Target Deconvolution. J. Proteome Res. 18, 4027–4037. 10.1021/acs.jproteome.9b00500.

8. Franken, H., Mathieson, T., Childs, D., Sweetman, G.M.A., Werner, T., Tögel, I., Doce, C., Gade, S., Bantscheff, M., Drewes, G., et al. (2015). Thermal proteome profiling for unbiased identification of direct and indirect drug targets using multiplexed quantitative mass spectrometry. Nat. Protoc. 10, 1567–1593. 10.1038/nprot.2015.101.

9. Demir, S., Wolff, G., Wieder, A., Maida, A., Bühler, L., Brune, M., Hautzinger, O., Feuchtinger, A., Poth, T., Szendroedi, J., et al. (2022). TSC22D4 interacts with Akt1 to regulate glucose metabolism. Sci. Adv. 8, eabo5555. 10.1126/sciadv.abo5555.

10. Ekim Üstünel, B., Friedrich, K., Maida, A., Wang, X., Krones-Herzig, A., Seibert, O., Sommerfeld, A., Jones, A., Sijmonsma, T.P., Sticht, C., et al. (2016). Control of diabetic hyperglycaemia and insulin resistance through TSC22D4. Nat. Commun. 7, 13267. 10.1038/ncomms13267.

11. Greenwald, W.W., Chiou, J., Yan, J., Qiu, Y., Dai, N., Wang, A., Nariai, N., Aylward, A., Han, J.Y., Kadakia, N., et al. (2019). Pancreatic islet chromatin accessibility and conformation reveals distal enhancer networks of type 2 diabetes risk. Nat. Commun. 10, 2078. 10.1038/s41467-019-09975-4.

12. Regué, L., Zhao, L., Ji, F., Wang, H., Avruch, J., and Dai, N. (2021). RNA m6A reader IMP2/IGF2BP2 promotes pancreatic β-cell proliferation and insulin secretion by enhancing PDX1 expression. Mol. Metab. 48, 101209. 10.1016/j.molmet.2021.101209.

13. Wolff, G., Sakurai, M., Mhamane, A., Troullinaki, M., Maida, A., Deligiannis, I.K., Yin, K., Weber, P., Morgenstern, J., Wieder, A., et al. (2022). Hepatocyte-specific activity of TSC22D4 triggers progressive NAFLD by impairing mitochondrial function. Mol. Metab. 60, 101487. 10.1016/j.molmet.2022.101487.

14. Van Vranken, J.G., Li, J., Mintseris, J., Wei, T.-Y., Sniezek, C.M., Gadzuk-Shea, M., Gygi, S.P., and Schweppe, D.K. (2024). Large-scale characterization of drug mechanism of action using proteome-wide thermal shift assays. Preprint at eLife Sciences Publications, Ltd, 10.7554/elife.95595.2 10.7554/elife.95595.2.

15. Pepelnjak, M., Velten, B., Näpflin, N., Von Rosen, T., Palmiero, U.C., Ko, J.H., Maynard, H.D., Arosio, P., Weber-Ban, E., De Souza, N., et al. (2024). In situ analysis of osmolyte mechanisms of proteome thermal stabilization. Nat. Chem. Biol. 10.1038/s41589-024-01568-7.

16. Noda, N., Jung, Y., Ado, G., Mizuhata, Y., Higuchi, M., Ogawa, T., Ishidate, F., Sato, S., Kurata, H., Tokitoh, N., et al. (2022). Glucose as a Protein-Condensing Cellular Solute. ACS Chem. Biol. 17, 567–575. 10.1021/acschembio.1c00849.

17. Gaetani, M., and Zubarev, R.A. (2023). Proteome Integral Solubility Alteration (PISA) for High-Throughput Ligand Target Deconvolution with Increased Statistical Significance and Reduced Sample Amount. Methods Mol. Biol. Clifton NJ 2554, 91–106. 10.1007/978-1-0716-2624-5_7.

18. Jackson, R.M., Griesel, B.A., Gurley, J.M., Szweda, L.I., and Olson, A.L. (2017). Glucose availability controls adipogenesis in mouse 3T3-L1 adipocytes via up-regulation of nicotinamide metabolism. J. Biol. Chem. 292, 18556–18564. 10.1074/jbc.M117.791970.

19. Jones, A., Friedrich, K., Rohm, M., Schäfer, M., Algire, C., Kulozik, P., Seibert, O., Müller-Decker, K., Sijmonsma, T., Strzoda, D., et al. (2013). TSC22D4 is a molecular output of hepatic wasting metabolism. EMBO Mol. Med. 5, 294–308. 10.1002/emmm.201201869.

20. Dai, N. (2020). The Diverse Functions of IMP2/IGF2BP2 in Metabolism. Trends Endocrinol. Metab. 31, 670–679. 10.1016/j.tem.2020.05.007.

21. Voight, B.F., Scott, L.J., Steinthorsdottir, V., Morris, A.P., Dina, C., Welch, R.P., Zeggini, E., Huth, C., Aulchenko, Y.S., Thorleifsson, G., et al. (2010). Twelve type 2 diabetes susceptibility loci identified through large-scale association analysis. Nat. Genet. 42, 579–589. 10.1038/ng.609.

22. Groenewoud, M.J., Dekker, J.M., Fritsche, A., Reiling, E., Nijpels, G., Heine, R.J., Maassen, J.A., Machicao, F., Schäfer, S.A., Häring, H.U., et al. (2008). Variants of CDKAL1 and IGF2BP2 affect first-phase insulin secretion during hyperglycaemic clamps. Diabetologia 51, 1659–1663. 10.1007/s00125-008-1083-z.

23. Tian, Y., Wan, N., Zhang, H., Shao, C., Ding, M., Bao, Q., Hu, H., Sun, H., Liu, C., Zhou, K., et al. (2023). Chemoproteomic mapping of the glycolytic targetome in cancer cells. Nat. Chem. Biol. 19, 1480–1491. 10.1038/s41589-023-01355-w.

24. Amnekar, R.V., Dite, T., Lis, P., Bell, S., Brown, F., Johnson, C., Wilkinson, S., Raggett, S., Dorward, M., Wightman, M., et al. (2024). NRBP1 pseudokinase binds to and activates the WNK pathway in response to osmotic stress. Preprint at Cold Spring Harbor Laboratory, 10.1101/2024.12.12.628181 10.1101/2024.12.12.628181.

25. Xiao, Y.-X., Lee, S.Y., Aguilera-Uribe, M., Samson, R., Au, A., Khanna, Y., Liu, Z., Cheng, R., Aulakh, K., Wei, J., et al. (2024). The TSC22D, WNK, and NRBP gene families exhibit functional buffering and evolved with Metazoa for cell volume regulation. Cell Rep. 43, 114417. 10.1016/j.celrep.2024.114417.

26. Brohée, S., and van Helden, J. (2006). Evaluation of clustering algorithms for protein-protein interaction networks. BMC Bioinformatics 7, 488. 10.1186/1471-2105-7-488.

27. Szklarczyk, D., Kirsch, R., Koutrouli, M., Nastou, K., Mehryary, F., Hachilif, R., Gable, A.L., Fang, T., Doncheva, N.T., Pyysalo, S., et al. (2023). The STRING database in 2023: protein-protein association networks and functional enrichment analyses for any sequenced genome of interest. Nucleic Acids Res. 51, D638–D646. 10.1093/nar/gkac1000.

28. Diabetes Genetics Initiative of Broad Institute of Harvard and MIT, Lund University, and Novartis Institutes of BioMedical Research, Saxena, R., Voight, B.F., Lyssenko, V., Burtt, N.P., de Bakker, P.I.W., Chen, H., Roix, J.J., Kathiresan, S., Hirschhorn, J.N., et al. (2007). Genome-wide association analysis identifies loci for type 2 diabetes and triglyceride levels. Science 316, 1331–1336. 10.1126/science.1142358.

29. Zeggini, E., Scott, L.J., Saxena, R., Voight, B.F., Marchini, J.L., Hu, T., de Bakker, P.I.W., Abecasis, G.R., Almgren, P., Andersen, G., et al. (2008). Meta-analysis of genome-wide association data and large-scale replication identifies additional susceptibility loci for type 2 diabetes. Nat. Genet. 40, 638–645. 10.1038/ng.120.

30. Scott, L.J., Mohlke, K.L., Bonnycastle, L.L., Willer, C.J., Li, Y., Duren, W.L., Erdos, M.R., Stringham, H.M., Chines, P.S., Jackson, A.U., et al. (2007). A Genome-Wide Association Study of Type 2 Diabetes in Finns Detects Multiple Susceptibility Variants. Science 316, 1341–1345. 10.1126/science.1142382.

31. Dai, N., Zhao, L., Wrighting, D., Krämer, D., Majithia, A., Wang, Y., Cracan, V., Borges-Rivera, D., Mootha, V.K., Nahrendorf, M., et al. (2015). IGF2BP2/IMP2 Deficient Mice Resist Obesity through enhanced translation of Ucp1 mRNA and other mRNAs encoding Mitochondrial Proteins. Cell Metab. 21, 609–621. 10.1016/j.cmet.2015.03.006.

32. Passaro, S., Corso, G., Wohlwend, J., Reveiz, M., Thaler, S., Somnath, V.R., Getz, N., Portnoi, T., Roy, J., Stark, H., et al. (2025). Boltz-2: Towards Accurate and Efficient Binding Affinity Prediction. Preprint at Cold Spring Harbor Laboratory, 10.1101/2025.06.14.659707 10.1101/2025.06.14.659707.

33. Burns, S.M., Vetere, A., Walpita, D., Dančík, V., Khodier, C., Perez, J., Clemons, P.A., Wagner, B.K., and Altshuler, D. (2015). High-throughput luminescent reporter of insulin secretion for discovering regulators of pancreatic Beta-cell function. Cell Metab. 21, 126–137. 10.1016/j.cmet.2014.12.010.

34. Yang, L., and Chen, W. (2023). Insulin secretion assays in an engineered MIN6 cell line. MethodsX 10, 102029. 10.1016/j.mex.2023.102029.

35. Miao, W., Porter, D.F., Siprashvili, Z., Ferguson, I.D., Ducoli, L., Nguyen, D.T., Ko, L.A., Lopez-Pajares, V., Srinivasan, S., Hong, A.W., et al. (2025). DDX50 cooperates with STAU1 to effect stabilization of pro-differentiation RNAs. Cell Rep. 44, 115174. 10.1016/j.celrep.2024.115174.

36. Varadi, M., Bertoni, D., Magana, P., Paramval, U., Pidruchna, I., Radhakrishnan, M., Tsenkov, M., Nair, S., Mirdita, M., Yeo, J., et al. (2024). AlphaFold Protein Structure Database in 2024: providing structure coverage for over 214 million protein sequences. Nucleic Acids Res. 52, D368–D375. 10.1093/nar/gkad1011.

37. Varadi, M., Anyango, S., Deshpande, M., Nair, S., Natassia, C., Yordanova, G., Yuan, D., Stroe, O., Wood, G., Laydon, A., et al. (2022). AlphaFold Protein Structure Database: massively expanding the structural coverage of protein-sequence space with high-accuracy models. Nucleic Acids Res. 50, D439–D444. 10.1093/nar/gkab1061.

38. Jumper, J., Evans, R., Pritzel, A., Green, T., Figurnov, M., Ronneberger, O., Tunyasuvunakool, K., Bates, R., Žídek, A., Potapenko, A., et al. (2021). Highly accurate protein structure prediction with AlphaFold. Nature 596, 583–589. 10.1038/s41586-021-03819-2.

39. Madeira, F., Madhusoodanan, N., Lee, J., Eusebi, A., Niewielska, A., Tivey, A.R.N., Lopez, R., and Butcher, S. (2024). The EMBL-EBI Job Dispatcher sequence analysis tools framework in 2024. Nucleic Acids Res. 52, W521–W525. 10.1093/nar/gkae241.

40. Waterhouse, A.M., Procter, J.B., Martin, D.M.A., Clamp, M., and Barton, G.J. (2009). Jalview Version 2—a multiple sequence alignment editor and analysis workbench. Bioinformatics 25, 1189–1191. 10.1093/bioinformatics/btp033.

41. Heinz, S., Benner, C., Spann, N., Bertolino, E., Lin, Y.C., Laslo, P., Cheng, J.X., Murre, C., Singh, H., and Glass, C.K. (2010). Simple combinations of lineage-determining transcription factors prime cis-regulatory elements required for macrophage and B cell identities. Mol. Cell 38, 576–589. 10.1016/j.molcel.2010.05.004.

42. ENCODE Project Consortium (2012). An integrated encyclopedia of DNA elements in the human genome. Nature 489, 57–74. 10.1038/nature11247.

43. Kagda, M.S., Lam, B., Litton, C., Small, C., Sloan, C.A., Spragins, E., Tanaka, F., Whaling, I., Gabdank, I., Youngworth, I., et al. (2025). Data navigation on the ENCODE portal. Nat. Commun. 16, 9592. 10.1038/s41467-025-64343-9.

44. Hitz, B.C., Lee, J.-W., Jolanki, O., Kagda, M.S., Graham, K., Sud, P., Gabdank, I., Seth Strattan, J., Sloan, C.A., Dreszer, T., et al. (2023). The ENCODE Uniform Analysis Pipelines. Preprint at Bioinformatics, 10.1101/2023.04.04.535623 10.1101/2023.04.04.535623.

45. Porter, D.F., Meyers, R.M., Miao, W., Reynolds, D.L., Hong, A.W., Yang, X., Srinivasan, S., Mondal, S., Siprashvili, Z., Fabo, T., et al. (2025). Disease-linked regulatory DNA variants and homeostatic transcription factors in epidermis. Nat. Commun. 16, 8387. 10.1038/s41467-025-63070-5.

46. Mendes, M.L., Fischer, L., Chen, Z.A., Barbon, M., O’Reilly, F.J., Giese, S.H., Bohlke-Schneider, M., Belsom, A., Dau, T., Combe, C.W., et al. (2019). An integrated workflow for crosslinking mass spectrometry. Mol. Syst. Biol. 15, e8994. 10.15252/msb.20198994.

47. Liu, N. (2019). Library Prep for CUT&RUN with NEBNext® Ultra^TM^ II DNA Library Prep Kit for Illumina® (E7645) v2. Preprint, 10.17504/protocols.io.bagaibse 10.17504/protocols.io.bagaibse.

48. Zarnegar, B.J., Webster, D.E., Lopez-Pajares, V., Vander Stoep Hunt, B., Qu, K., Yan, K.J., Berk, D.R., Sen, G.L., and Khavari, P.A. (2012). Genomic Profiling of a Human Organotypic Model of AEC Syndrome Reveals ZNF750 as an Essential Downstream Target of Mutant TP63. Am. J. Hum. Genet. 91, 435–443. 10.1016/j.ajhg.2012.07.007.

49. Cox, J., and Mann, M. (2008). MaxQuant enables high peptide identification rates, individualized p.p.b.-range mass accuracies and proteome-wide protein quantification. Nat. Biotechnol. 26, 1367–1372. 10.1038/nbt.1511.

50. Ikaga, R., Namekata, I., Kotiadis, V.N., Ogawa, H., Duchen, M.R., Tanaka, H., and Iida-Tanaka, N. (2015). Knockdown of aquaporin-8 induces mitochondrial dysfunction in 3T3-L1 cells. Biochem. Biophys. Rep. 4, 187–195. 10.1016/j.bbrep.2015.09.009.

51. Wohlwend, J., Corso, G., Passaro, S., Getz, N., Reveiz, M., Leidal, K., Swiderski, W., Atkinson, L., Portnoi, T., Chinn, I., et al. (2024). Boltz-1 Democratizing Biomolecular Interaction Modeling. Preprint at Biophysics, 10.1101/2024.11.19.624167 10.1101/2024.11.19.624167.

52. Jumper, J., Evans, R., Pritzel, A., Green, T., Figurnov, M., Ronneberger, O., Tunyasuvunakool, K., Bates, R., Žídek, A., Potapenko, A., et al. (2021). Highly accurate protein structure prediction with AlphaFold. Nature 596, 583–589. 10.1038/s41586-021-03819-2.

53. Varadi, M., Anyango, S., Deshpande, M., Nair, S., Natassia, C., Yordanova, G., Yuan, D., Stroe, O., Wood, G., Laydon, A., et al. (2022). AlphaFold Protein Structure Database: massively expanding the structural coverage of protein-sequence space with high-accuracy models. Nucleic Acids Res. 50, D439–D444. 10.1093/nar/gkab1061.

54. Porter, D.F., Miao, W., Yang, X., Goda, G.A., Ji, A.L., Donohue, L.K.H., Aleman, M.M., Dominguez, D., and Khavari, P.A. (2021). easyCLIP analysis of RNA-protein interactions incorporating absolute quantification. Nat. Commun. 12, 1569. 10.1038/s41467-021-21623-4.

55. The UniProt Consortium, Bateman, A., Martin, M.-J., Orchard, S., Magrane, M., Adesina, A., Ahmad, S., Bowler-Barnett, E.H., Bye-A-Jee, H., Carpentier, D., et al. (2025). UniProt: the Universal Protein Knowledgebase in 2025. Nucleic Acids Res. 53, D609–D617. 10.1093/nar/gkae1010.

56. Katoh, K., and Standley, D.M. (2013). MAFFT multiple sequence alignment software version 7: improvements in performance and usability. Mol. Biol. Evol. 30, 772–780. 10.1093/molbev/mst010.

57. Shannon, C.E. (1948). A Mathematical Theory of Communication. Bell Syst. Tech. J. 27, 379–423. 10.1002/j.1538-7305.1948.tb01338.x.

